# A novel matrix multiplication framework for modeling genotype-by-environment interaction in genomic prediction

**DOI:** 10.64898/2026.05.11.724414

**Authors:** Osval A. Montesinos-López, Abelardo Montesinos-López, José Cricelio Montesinos-Lopez, José Crossa, Susanne Dreisigacker, Carlos M. Hernández Suárez, Rodomiro Ortiz

**Author notes:** Corresponding authors: Abelardo. Montesinos-López, and Rodomiro Ortiz.

## Abstract

Accurate modeling of genotype-by-environment (G×E) interaction is critical for genomic prediction in plant breeding but remains challenging due to complex interaction structures. Conventional models often use the Hadamard product of genotype and environment covariance matrices to capture joint similarity, which may not fully represent G×E complexity. Here we propose a novel framework that derives covariance structures from the matrix multiplication of genotype and environment kernels, decomposing these into symmetric components incorporated as random effects in mixed models. Evaluated for 11 wheat and rice multi-environment datasets and across, this approach consistently outperformed the traditional Hadamard-based model, improving prediction accuracy by up to 13.2% in Pearson’s correlation and enhancing top-selection accuracy. Combining both methods yielded the highest performance, indicating complementary information capture. This framework offers a flexible, interpretable, and computationally feasible extension for modeling G×E interaction, potentially enhancing genomic selection effectiveness under diverse environmental conditions.

## INTRODUCTION

Genotype-by-environment (G×E) interaction, which refers to the phenomenon where different genotypes respond differently to varying environmental conditions, is a central challenge in quantitative genetics and plant breeding. This interaction determines how genetic potential is expressed across diverse environments, often resulting in changes in genotype rankings and performance across locations and years. Such variability complicates selection decisions and can reduce the effectiveness of breeding programs if not properly modeled. Therefore, accurately capturing G×E interaction is essential for improving the prediction of genetic values, identifying broadly adapted genotypes, and enhancing genetic gain under heterogeneous and changing environments (Cooper et al. 1999; Crossa 1990; Crossa et al. 2017).

The statistical modeling of G×E interaction has evolved substantially over the last century. Early approaches, such as the analysis of variance, partitioned phenotypic variation into genetic, environmental, and interaction components (Fisher 1925), but often regarded G×E primarily as a nuisance term. Regression-based stability models, including those of Finlay and Wilkinson (1963) and Eberhart and Russell (1966), introduced quantitative methods to characterize genotype responses across environmental gradients. Crossa (1990) expanded on these foundations by emphasizing the integration of stability analysis with predictive accuracy. Later, researchers such as Cornelius et al. (1992) and Cornelius and Crossa (1999) advanced the field by clarifying the importance of structured covariance models and multiplicative interaction components, which underpin many modern approaches to modeling G×E.

Subsequent developments introduced multivariate and mixed model methods, such as additive main effects and multiplicative interaction (AMMI) models (Gauch 1988), which decompose genotype and environment effects into additive and interaction components, and genotype main effect plus G×E interaction (GGE) biplots (Yan et al. 2000), which visually represent genotype and environment performance. These approaches offer both interpretability and predictive capacity and remain widely used in the analysis of multi-environment trials.

With the advent of genomic selection (Meuwissen et al. 2001), the focus shifted toward predictive modeling, aiming to estimate the performance of untested genotypes using genome-wide marker data. However, many early genomic prediction models either ignored G×E interactions altogether or incorporated them using oversimplified assumptions, leading to reduced prediction accuracy in multi-environment trials.

To address this limitation, linear mixed models incorporating structured covariance matrices were developed (Crossa et al. 2010; Burgueño et al. 2012). These developments were further consolidated by the integration of genomic prediction and G×E modeling frameworks, which combine marker-based relationships with environmental information to improve prediction accuracy in multi-environment trials (Crossa et al. 2017).

A widely adopted strategy for modeling G×E interaction in genomic prediction is based on the Hadamard (elementwise) product of genotype and environment covariance matrices. This formulation captures interaction effects through joint similarity: interaction strength is large when genotypes are genetically similar, and environments are environmentally similar. The approach is attractive due to its biological interpretability, statistical parsimony, and computational efficiency, and has been successfully applied in multi-environment genomic prediction models (Jarquín et al. 2014; Cuevas et al. 2017). Nevertheless, the Hadamard product represents only one way to combine genotype and environmental information and implicitly assumes that interaction effects can be fully characterized by joint similarity structures.

Recent developments in kernel methods and multi-source data integration suggest that alternative combinations of covariance structures may capture additional aspects of complex biological interactions (Cuevas et al., 2025). For example, Cuevas et al. (2025) proposed a framework based on matrix multiplication for integrating pedigree and genomic information, deriving two complementary covariance matrices that are modeled as separate random effects. This approach has demonstrated improved predictive performance when integrating multiple sources of biological information, such as genomics, transcriptomics, metabolomics, and phenomics (Montesinos-López et al. 2025a,b). Given that G×E interaction also involves the integration of genetic and environmental information, adapting this matrix multiplication framework to model G×E interaction could potentially capture more complex and structured relationships than the conventional Hadamard approach. However, despite its success in multi-source integration, this framework has not yet been explored for modeling G×E interaction.

The key contribution of this study is the adaptation of the framework of Cuevas et al. (2025) to explicitly model G×E interaction. While this framework was originally developed for integrating multiple sources of biological information, we demonstrate that it can be extended to G×E modeling, resulting in improved prediction accuracy. Unlike the conventional Hadamard product, which captures interaction through joint similarity between genotypes and environments, the proposed framework extracts additional structured information from the matrix product of genotype and environment kernels. This matrix product allows the model to capture more complex, potentially directional, and higher-order relationships that are not accessible through elementwise operations. Importantly, these matrix-product-based kernels provide information that is complementary to, rather than redundant with, the Hadamard-based formulation.

Building on this idea, we extend the framework of Cuevas et al. (2025) to explicitly model G×E interaction in genomic prediction. Specifically, we construct two covariance matrices by decomposing the matrix product of genotype and environment kernels into its upper and lower triangle components and incorporate these as random effects in linear mixed models. This approach results in a family of models that either replace or augment the conventional Hadamard-based interaction term. We hypothesize that combining both the Hadamard and matrix-product-based approaches will provide a more complete representation of G×E structure.

To evaluate this hypothesis, we analyze 11 independent datasets from wheat and rice multi-environment trials. Our results show that models based on matrix-product-derived kernels improve prediction accuracy relative to the conventional Hadamard approach, and that the combination of both strategies yields the highest predictive performance. These findings suggest that G×E interaction is inherently multi-structured and cannot be fully captured by a single covariance formulation.

Overall, this work provides a new perspective on modeling G×E interaction in genomic prediction by introducing complementary kernel structures that enhance predictive accuracy beyond what is achievable with conventional methods. The proposed framework is general, flexible, and readily implementable within standard mixed-model software, offering a practical and effective tool for modern plant breeding programs operating under complex environmental conditions.

## MATERIALS AND METHODS

### Datasets

#### Datasets wheat EYT_1, EYT_2, and EYT_3

The datasets used in this study correspond to elite yield trial (EYT) cycles from the Global Wheat Program of the International Maize and Wheat Improvement Center (CIMMYT). Specifically, they include the 2013–2014, 2014–2015, and 2015–2016 cycles, hereafter referred to as EYT_1, EYT_2, and EYT_3, respectively. These trials were conducted across four cropping seasons, each evaluated in four or five environments.

The study included 776 lines in EYT_1,775 in EYT_2, and 964 in EYT_3. Trials were conducted using an alpha lattice design, with 39 trials in total. Each trial consisted of 28 lines and two checks, arranged into six blocks with three replications. Across the datasets, several traits were evaluated depending on the environment and lines. For consistency, four traits were analyzed for each line in every environment: (i) days to heading, defined as the number of days from germination until 50% spike emergence, (ii) days to maturity, defined as the number of days from germination until 50% physiological maturity, or when 50% of spikes had lost their green color, (iii) plant height, and (iv) grain yield (GY). Further details regarding the experimental design and the computation of best linear unbiased estimates are provided in Juliana et al. (2018).

Lines from EYT_1 were evaluated in four environments: bed planting with five irrigations (Bed5IR), early heat (EHT), flat planting with five irrigations (Flat5IR), and late heat (LHT). EYT_2 was assessed in five environments: bed planting with two irrigations (Bed2IR), Bed5IR, EHT, Flat5IR, and LHT. Similarly, EYT_3 was evaluated in five environments: Bed2IR, Bed5IR, Flat5IR, flat planting with drip irrigation (FlatDrip), and LHT.

Genome-wide markers for the 2,515 lines (776 + 775 + 964) were obtained through genotyping-by-sequencing (GBS; Elshire et al., 2011; Poland et al., 2012)) at Kansas State University using the Illumina HiSeq2500 platform. From an initial set of 34,900 markers, 2,038 were retained after quality filtering. Missing marker data were imputed with LinkImpute (Money et al., 2015), implemented in TASSEL v5 (Bradbury et al., 2007). Lines with more than 50% missing data were excluded, resulting in a final dataset of 2,515 lines across the three EYT cycles.

#### Datasets 4. Indica

The Indica dataset (Monteverde et al., 2019) is a rice resource that includes phenotypic records for four traits: Percentage of Head Rice Recovery (PHR, g), measured as the weight of intact milled kernels from a 100 g rough rice sample; Grain Yield (GY, kg/ha) of paddy rice; Plant Height (PH, cm), recorded from the soil surface to the tip of the flag leaf; and Grain Chalkiness (GC, %), quantified as the proportion of chalky kernels in a 50 g subsample of total milled rice. These traits were collected across three growth stages—vegetative, reproductive, and maturation—over three environments (2010, 2011, and 2012). In each environment, 327 genotypes were evaluated, resulting in a total of 981 observations, with each genotype tested once per environment. After quality control, 16,383 SNP markers were retained and coded as 0, 1, or 2. Further details are provided in **Table 1** and in Monteverde et al. (2019).

**Table 1.**
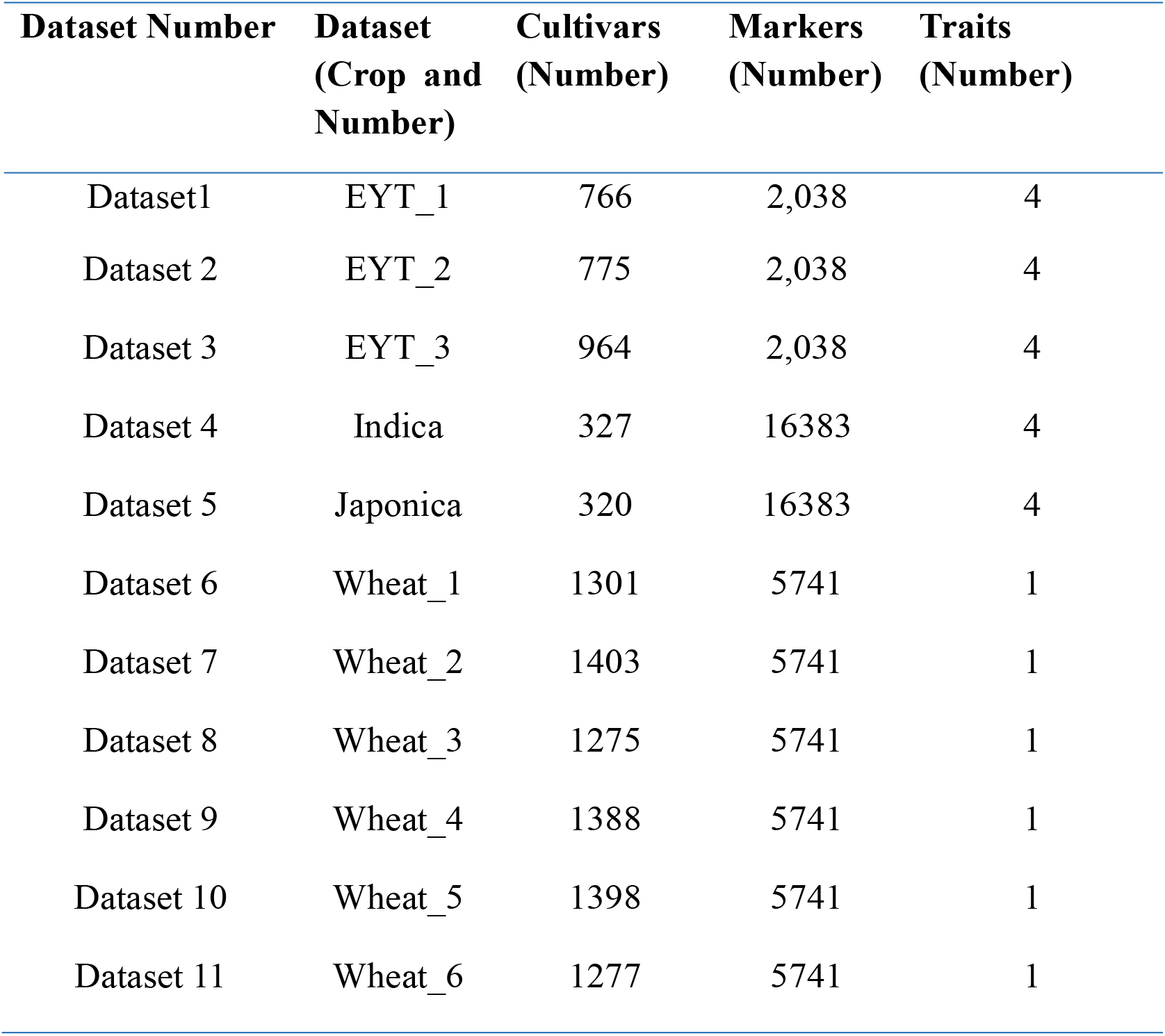
Description of the datasets.

| **Dataset Number** | **Dataset**<br>**(Crop and**<br>**Number)** | **Cultivars**<br>**(Number)** | **Markers**<br>**(Number)** | **Traits**<br>**(Number)** |
| --- | --- | --- | --- | --- |
| Dataset1 | EYT_1 | 766 | 2,038 | 4 |
| Dataset 2 | EYT_2 | 775 | 2,038 | 4 |
| Dataset 3 | EYT_3 | 964 | 2,038 | 4 |
| Dataset 4 | Indica | 327 | 16383 | 4 |
| Dataset 5 | Japonica | 320 | 16383 | 4 |
| Dataset 6 | Wheat_1 | 1301 | 5741 | 1 |
| Dataset 7 | Wheat_2 | 1403 | 5741 | 1 |
| Dataset 8 | Wheat_3 | 1275 | 5741 | 1 |
| Dataset 9 | Wheat_4 | 1388 | 5741 | 1 |
| Dataset 10 | Wheat_5 | 1398 | 5741 | 1 |
| Dataset 11 | Wheat_6 | 1277 | 5741 | 1 |

#### Datasets 5. Japonica

The Japonica dataset consists of 320 genotypes from the tropical rice Japonica population. It was evaluated for the same four traits as the Indica dataset—Plant Height (PH), Percentage of Head Rice Recovery (PHR), Grain Yield (GY), and Grain Chalkiness (GC)—across five environments from 2009 to 2013. In total, 1,051 observations were recorded, although the dataset is unbalanced. Each genotype was also genotyped with 16,383 SNP markers retained after quality control, coded as 0, 1, or 2. Additional details are available in Monteverde et al. (2019).

#### Datasets 6-11

Six data sets of spring wheat lines tested for grain yield from CIMMYT’s first-year yield trials (YT), as described in Ibba et al. (2020), were used in this study:

- Wheat_1 (YT2013-14/EYT2014-15): 1,301 lines from the 2013–14 YT and 472 lines from the 2014–15 EYT.
- Wheat_2 (YT2014-15/EYT2015-16): 1,337 lines from the 2014–15 YT and 596 lines from the 2015–16 EYT.
- Wheat_3 (YT2015-16/EYT2016-17): 1,161 lines from the 2015–16 YT and 556 lines from the 2016–17 EYT.
- Wheat_4 (YT2016-17/EYT2017-18): 1,372 lines from the 2016–17 YT and 567 lines from the 2017–18 EYT.
- Wheat_5 (YT2017-18/EYT2018-19): 1,386 lines from the 2017–18 YT and 509 lines from the 2018–19 EYT.
- Wheat_6 (YT2018-19/EYT2019-20): 1,276 lines from the 2018–19 YT and 124 lines from the 2019–20 EYT.

All lines were genotyped using genotyping-by-sequencing (GBS). Marker polymorphisms were called with the TASSEL v5 GBS pipeline, applying a minor allele frequency (MAF) threshold of 0.01 for SNP discovery. A total of 6,075,743 unique tags were aligned to the wheat reference genome (RefSeq v.1.0; IWGSC 2018), achieving an alignment rate of 63.98%. After filtering SNPs for >80% homozygosity, *p* < 0.001 based on Fisher’s exact test, and χ^2^ values below the critical threshold of 9.2, 78,606 GBS markers passed at least one of these criteria. Additional filtering retained markers with <50% missing data, MAF>0.05, and <5% heterozygosity. Missing data were imputed using the expectation–maximization algorithm implemented in the R package rrBLUP (Endelman, 2011).

**Table 1** outlines the general characteristics of the 11 public datasets used in this study. Nine datasets correspond to wheat trials, and two correspond to rice sets (Indica and Japonica). The public repository available at https://github.com/osval78/New_GE_Framework contains the phenotypic and genomic data for all datasets, along with the codes used for implementing the proposed methodology.

### Statistical models

Theory on models is in the **Appendix Theory**

#### Model A

The predictive response model is defined as:

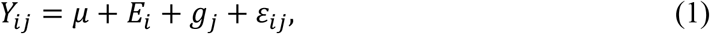

where *Y*_*ij*_ denotes the response variable measured for genotype *j* in environment *i*; *μ* denotes the general mean or intercept; *E*_*i*_, *i* = 1, …, *I*, denotes the random effects of environments, distributed as 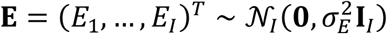, with **I** being the identity matrix of order *I; g*_*j*_, *j* = 1, …, *J*, are the random effects of lines; and *ϵ*_*ij*_ denotes the random error terms, assumed normally distributed with mean 0 and variance *σ*^2^. Furthermore, it is assumed that 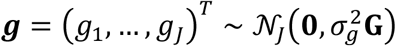, where **G** denotes the linear kernel computed as 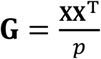. This matrix is known as the genomic relationship matrix (VanRaden, 2008) in plant and animal breeding, since it is computed from a standardized matrix of SNPs (**X**), coded as 0, 1, and 2, of dimension *J* × *p*. The matrix **G** is considered known before the modelling process.

Let **y** be the stacked vector of all observations *Y*_*ij*_ (of length n = IJ). A common matrix formulation of model (1) is:

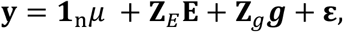

where **1**_*n*_ is the *n* × 1 vector of ones, **Z**_*E*_ is the *n* × *I* incidence matrix linking observations to environment, **Z**_*g*_ is the *n* × *J* incidence matrix linking observations to genotypes (lines), and **ε** is a vector of residuals distributed as **ε** ~ *N*_*n*_(0, *σ*^2^**I**_*n*_).

#### Model B

Model B extends model A by including G×E interaction effects:

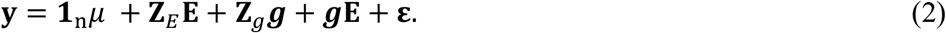

All terms are defined as in model A, except the interaction term ***g*E** which denotes the G×E interaction effects, ***g*E** = (*gE*_11_, …, *gE*_1*J*_, …, *gE*_*IJ*_)^*T*^, that are modeled as a multivariate normal distribution 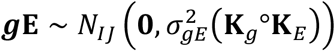, where 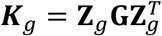, with ***Z***_*g*_ the *n* × *J* incidence matrix for additive genetic effects, the variance component 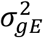 corresponds to the G×E interaction, 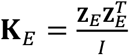 represents the linear kernel of the environments, with **Z**_*E*_ is the *n* × *I* incidence matrix for the environmental effects. ° denotes the Hadamard (elementwise) product. The Hadamard product **K**_*g*_*°***K**_*E*_ captures the variance-covariance structure of the **G×E** interaction effects because: (a) 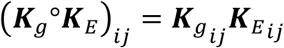 is large only when both genetic similarity 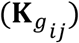 and environmental similarity 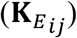 are large, where 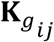 denotes (*i, j*) the entry of a matrix **K**_*g*_, and b) it captures joint similarity, modeling effects that are specific to particular genetic-environmental combinations. Since both **K**_*g*_ and **K**_*E*_ are symmetric and positive semi-definite (PSD) variance-covariance matrices (see proof in **Appendix theory**) their Hadamard product **(K**_*g*_*°***K**_*E*_) is also PSD, ensuring a valid variance-covariance matrix.

#### Model C

Model C extends Model A by incorporating an alternative structure for the G×E interaction:

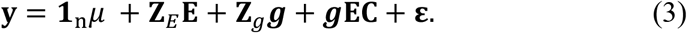

All components match those in Model A, except for the term ***g*EC** that denotes the G×E interaction effects. Let **gEC** = (*gEC*_11_, …, *gEC*_1*J*_, …, *gEC*_*IJ*_)^*T*^. These interaction effects are modeled as follow a multivariate normal distribution, 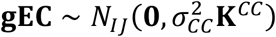, where 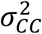 is the variance component associated with this interaction, and **K**^*CC*^ is a matrix derived from the matrix multiplication product of the environmental **K**_*E*_ and **K**_*g*_ genetic kernels. Let **M** = **K**_*E*_**K**_*g*_, then decomposes **M** into three mutually exclusive parts:

- **Upper-triangular off-diagonal part (U)** 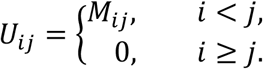
- **Diagonal part (D)** 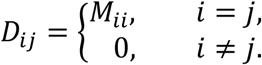
- **Lower-triangular off-diagonal part (L)**

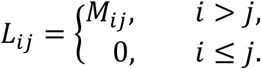

Thus, **M** = **L** + **D** + **U**. Following Cuevas et al. (2025), the covariance matrix for Model C is constructed as **K**^*CC*^ *=* **U** + **D** + **U**^*T*^. This construction preserves the diagonal of **M** and symmetrizes the upper-triangular information, producing a valid symmetric covariance matrix (Cuevas et al., 2025).

#### Model D

Model D extends model A by incorporating an alternative structure for the G×E interaction:

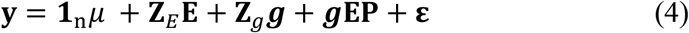

All components are as defined previously, except for the term ***g*EP**, which denotes an alternative G×E interaction effect. Let ***g*EP** = (*gEP*_11_, …, *gEP*_1*J*_, …, *gEP*_*IJ*_)^*T*^. These effects are modeled as following a multivariate normal distribution 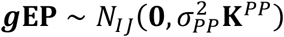, where 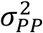 is the variance component associated with the random effect ***g*EP**.

Using the same decomposition of the matrix multiplication product **M** = **K**_*E*_**K**_*g*_ into **L, D**, and **U**, the covariance matrix for Model D is based on the lower-triangular component:

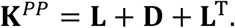

This formulation preserves the diagonal of **M** and symmetrizes the lower-triangular information, yielding a second valid covariance matrix that captures complementary structure to **K**^*PP*^.

#### Model E

Model E extends the previous models by incorporating both upper and lower triangular interaction structures:

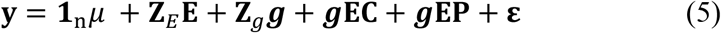

All terms correspond to those specified in the preceding models. As shown in equation (5), this model incorporates the interaction terms defined in equations (3) and (4), aiming to fully capture the complex interaction accounted by the upper and lower triangular matrices derived from the matrix product of the ***K***_*E*_ and ***K***_*g*_ linear kernels.

#### Model F

Model F incorporates all previously defined components of the G×E interaction:

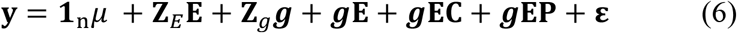

All terms remain consistent with those used in the earlier models. The motivation for Model F is that the two approaches for modelling the G×E interaction: the Hadamard-based kernel (Model B), and the matrix-multiplication-based kernels (Models C and D) capture different aspects of the underlying interaction structure: the Hadamard kernel **K**_*g*_*°***K**_*E*_ models interaction effects driven by joint similarity between genotypes and environments; the triangular-based kernels **K**^*CC*^ and **K**^*PP*^ capture directional or asymmetric patterns emerging from the matrix multiplication product **K**_*E*_**K**_*g*_, which may reflect more complex or structured interaction behaviors.

All models were implemented in the R statistical software (R Core Team, 2025) using the BGLR package (Pérez and de los Campos, 2014). Since this is a Bayesian statistical software, we used 10,000 iterations, with a burning of 2500 and a thinning of 5.

Models A and B correspond to conventional specifications: Model A includes only main effects, whereas Model B incorporates both main effects and G×E interactions modeled via the Hadamard product. In contrast, Models C through F represent the proposed extensions, obtained by adapting the matrix-product-based framework of Cuevas et al. (2025) to the specific context of G×E interaction. Model C extends Model A by including an additional random effect constructed from the upper triangular component of the matrix product between ***K***_*E*_and ***K***_*g*_. Similarly, Model D extends Model A by incorporating a random effect based on the lower triangular component of the same matrix product. Model E further augments Model A by including two random effects, derived from both the upper and lower triangular components of the matrix product between ***K***_*E*_and ***K***_*g*_. Finally, Model F builds upon Model B by adding these two random effects, thereby combining the conventional Hadamard-based interaction term with those derived from matrix multiplication. From these formulations, we hypothesize that Model F will outperform Model B, as it not only retains the conventional G×E interaction term based on the Hadamard product but also incorporates additional structure through the two random effects derived from matrix multiplication. Moreover, we expect Model F to achieve higher prediction accuracy than Models C, D, and E, since these models rely exclusively on the matrix-product-based components—either partially (Models C and D) or fully (Model E)—while omitting the conventional interaction term Compliance with the assumptions of the proposed models is provided in the **Appendix Theory**.

### Cross-validation and evaluation metrics

To evaluate and compare the predictive performance of the models, we employed a cross-validation strategy that simulated the scenario of untested lines in tested environments (ULTE). This approach involved five random partitions of the full dataset, with each partition using 50% of the observations for training and the remaining 50% for testing (Alemu et al., 2006; Burgueño et al., 2012). Predictive performance was assessed using the proportion of test items whose predicted top 10% lines match the true top 10%(PM_10) and the Average Pearson Correlation (COR), both calculated for each testing set within every partition. The overall prediction accuracy for each dataset was reported as the average PM_10 and COR values across all environments/traits and partitions.

## RESULTS

We provide the results for data sets EYT_1, Indica, Japonica, and Wheat_1 in Tables 2-5, Figures 1-4, and Appendix A, Table 2A. Results across the 11 data sets under study are in Table 6, Figure 5, and Appendix A, Table 1A. Results for data sets EYT_2, EYT_3, and Wheat 2-6 are given in Appendix A Table A1, and Appendix B, Table B1.

**Table 2.** Proportion of total variance, across traits in data set EYT_1, explained for each variance component in each of the evaluated models. VGE_A = VGE + VGEC + VGEP, which denotes the total proportion of variance explained by the G×E interaction. 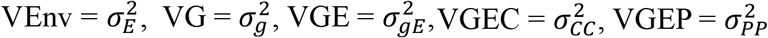, and VE = *σ*^2^.

| Dataset | Methods | VEnv | VG | VGE | VGEC | VGEP | Ve | VGE A |
| --- | --- | --- | --- | --- | --- | --- | --- | --- |
| EYT_1 | A | 0.719 | 0.188 | - | - | - | 0.093 | 0.000 |
| EYT_1 | B | 0.695 | 0.208 | 0.034 | - | - | 0.064 | 0.034 |
| EYT_1 | C | 0.711 | 0.190 | - | 0.011 | - | 0.088 | 0.011 |
| EYT_1 | D | 0.712 | 0.192 | - | - | 0.009 | 0.087 | 0.009 |
| EYT_1 | E | 0.706 | 0.195 | - | 0.014 | 0.011 | 0.074 | 0.025 |
| EYT_1 | F | 0.677 | 0.218 | 0.028 | 0.010 | 0.008 | 0.059 | 0.046 |

**Figure 1.**
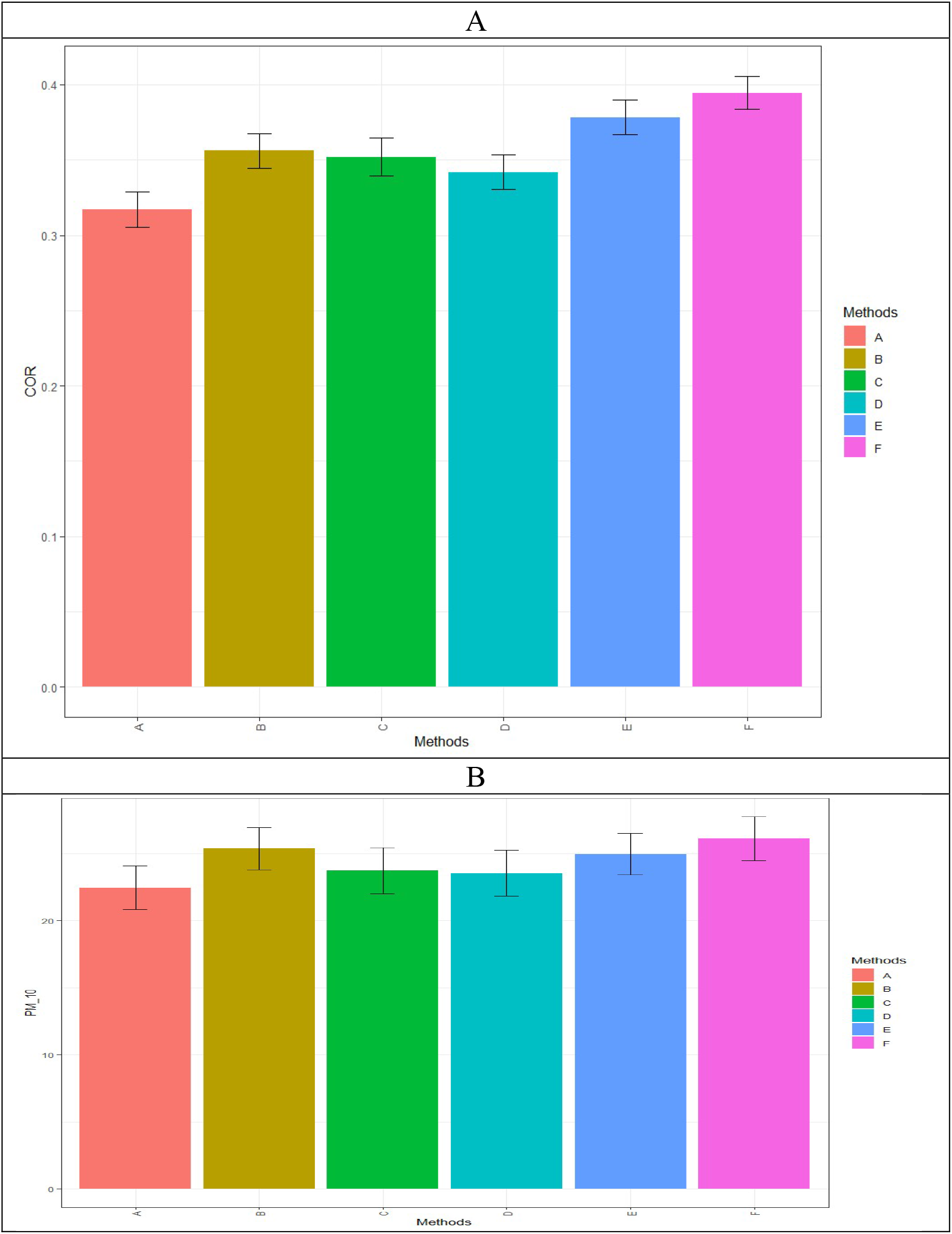
Prediction performance of the six evaluated methods on the EYT_1 dataset, measured by average Pearson’s correlation (COR; A) and the proportion of test items whose predicted top 10% lines match the true top 10% (PM_10, B).

### EYT_1 dataset

**Table 2** shows that, across all models for this dataset, the environment component (VEnv = 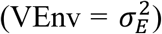) explained the largest proportion of explained variance, followed by the genotype component (VG = 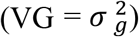). Among the G×E components (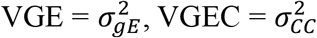, and VGEP = 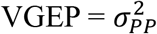), only model F accounted for a substantially larger proportion (a 35% increase) of the total variability explained (VGE_A = VGE + VGEC + VGEP) compared with model B, which included the conventional G×E interaction term.

Model F achieved the highest prediction performance in terms of COR (COR = 0.395), followed by model E (COR = 0.378), which was 4.5% lower than model F (Figure 1A). Model A had the lowest performance (COR = 0.317; **Figure 1A**), representing a 24.6% decrease compared to model F, while model D ranked second lowest with a COR of 0.342 (15.5% lower than model F).

The PM_10 model F exhibited the highest predictive performance (PM_10 = 26.122), followed closely by model B (PM_10 = 25.369), which was 3% lower than that of model F (Figure 1B). Model A had the lowest performance (PM_10 = 22.452; **Figure 1B**), a 16.3% reduction relative to model F, while model D was the second lowest (PM_10 = 23.452; 1% lower than model F). More details of the prediction performance for this data set are given in Table A2, Appendix A.

### Indica dataset

**Table 3** shows that, across all models, the genotype component contributed the largest share of explained variance, followed by the error component. Among the G×E components (VGE, VGEC, and VGEP), only model F captured a greater proportion of the total variability (a 17.1% increase) compared with model B, which used only the conventional G×E interaction term.

**Table 3.** Proportion of total variance, across traits in data set Indica, explained for each variance component in each of the evaluated models. VGE_A = VGE + VGEC + VGEP, which denotes the total proportion of variance explained by the G×E interaction. 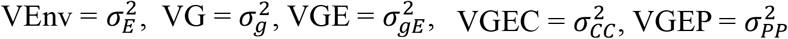, and VE = *σ*^2^.

| Dataset | Methods | VEnv | VG | VGE | VGEC | VGEP | Ve | VGE A |
| --- | --- | --- | --- | --- | --- | --- | --- | --- |
| Indica | A | 0.056 | 0.493 | - | - | - | 0.450 | 0.000 |
| Indica | B | 0.048 | 0.541 | 0.105 | - | - | 0.306 | 0.105 |
| Indica | C | 0.056 | 0.525 | - | 0.027 | - | 0.392 | 0.027 |
| Indica | D | 0.047 | 0.538 | - | - | 0.023 | 0.392 | 0.023 |
| Indica | E | 0.051 | 0.552 | - | 0.031 | 0.030 | 0.337 | 0.061 |
| Indica | F | 0.042 | 0.566 | 0.088 | 0.022 | 0.013 | 0.269 | 0.123 |

For COR, model F achieved the highest predictive accuracy (COR = 0.405), followed by model E (COR = 0.399), which was 1.5% lower than that of model F (**Figure 2A**). Model A had the lowest performance (COR = 0.359; **Figure 2A**), a 12.8% decline relative to model F, while model C ranked second lowest (COR of 0.372, lower than model F).

**Figure 2.**
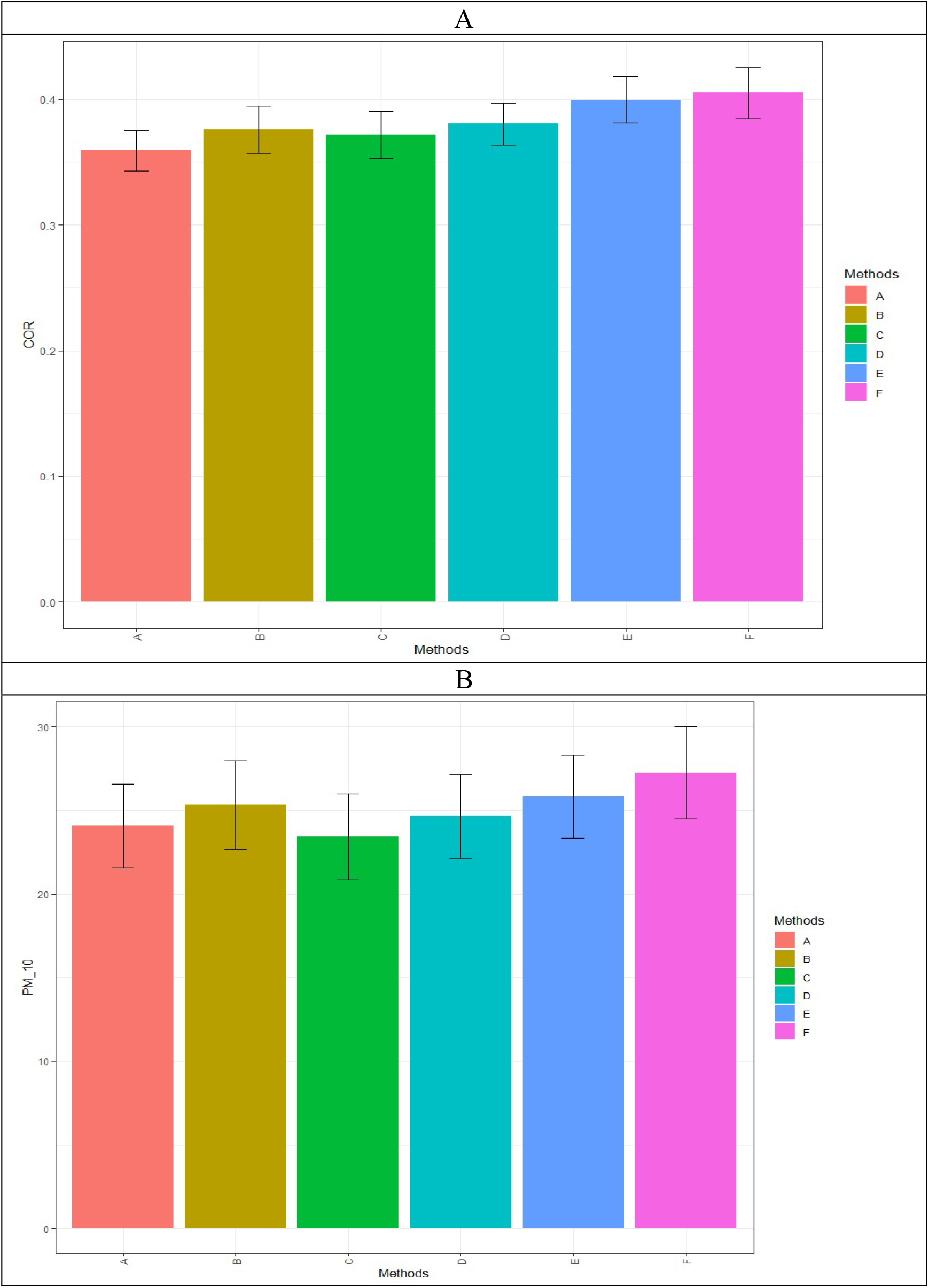
Prediction performance of the six evaluated methods on the Indica dataset, measured by average Pearson’s correlation (COR; A) and the proportion of test items whose predicted top 10% lines match the true top 10% (PM_10, B).

For PM_10, model F delivered the highest predictive outcome (PM_10 = 27.255), followed closely by model E (PM_10 = 25.833), which was 5.5% below that of model F (**Figure 2B**). Model C had the lowest result (PM_10 = 23.431; **Figure 2B**), a 16.3% decrease compared to model F. Model A followed as the second lowest (PM_10 of 24.069; 13.2% lower than model F). More details of the prediction performance for this data set are given in Table A2, Appendix A.

### Japonica dataset

The environment component accounted for the largest proportion of explained variance, followed by the genotype component across all models (**Table 4**). Among the G×E components (VGE, VGEC, and VGEP), only model F explained a higher share of the total variability (a 12.162% increase) compared to model B, which included only the standard G×E interaction term.

**Table 4.**
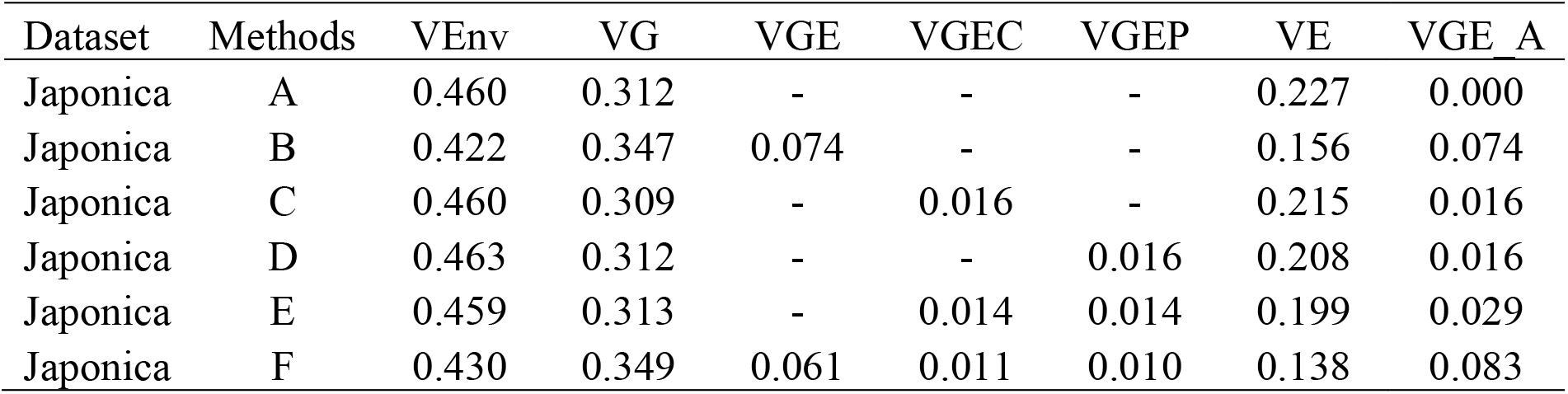
Proportion of total variances, in dataset Japonica, explained for each variance component in each of the evaluated models. VGE_A = VGE + VGEC + VGEP, which denotes the total proportion of variance explained by the G×E interaction. 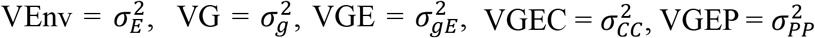, and VE = *σ*^2^.

For COR, model F achieved the highest predictive accuracy (COR = 0.421), followed by model E (COR = 0.413), which was 1.9% lower than model F (**Figure 3A**). Model A had the lowest performance (COR = 0.397; **Figure 3A**), a 6.0% decrease compared to model F, while models C and D tied for the second lowest (COR = 0.404, representing a 4.2% reduction relative to model F.

**Figure 3.**
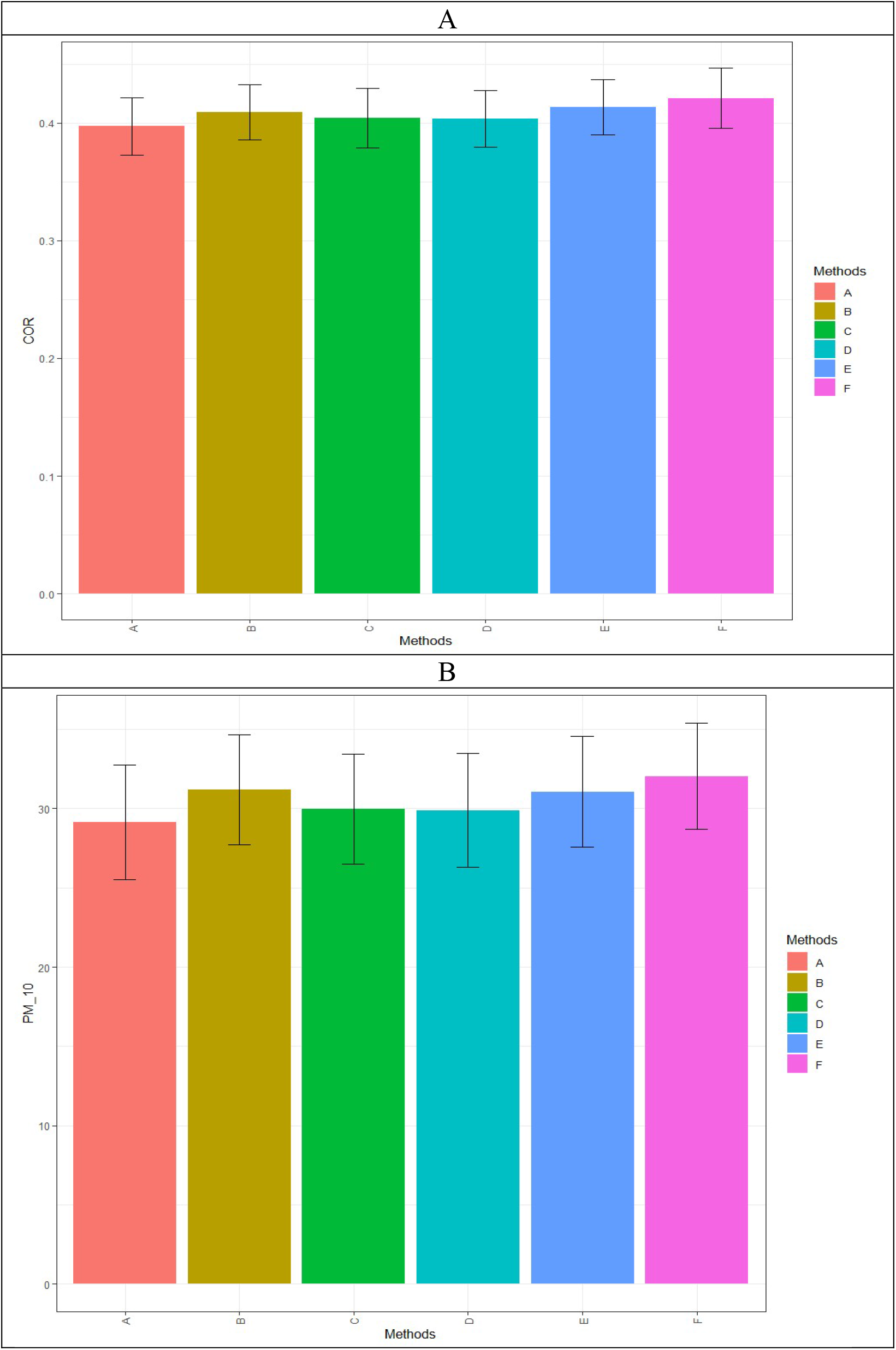
Prediction performance of the six evaluated methods on the Japonica dataset, measured by average Pearson’s correlation (COR; A) and the proportion of test items whose predicted top 10% lines match the true top 10% (PM_10, B).

For COR, model F achieved the highest predictive accuracy (COR = 0.421), followed by model E (COR = 0.413), which was 1.9% lower than model F (Figure 3A). Model A had the lowest performance (COR = 0.397; **Figure 3A**), a 6.0% decrease compared to model F, while models C and D tied for the second lowest (COR = 0.404; 4.2% lower than model F).

Regarding PM_10, model F produced the highest predictive value (PM_10 = 32.038), followed closely by model B (PM_10 = 31.188), which was 2.7% lower than that of model F (Figure 3B). In contrast, model A reported the weakest outcome (PM_10 = 29.138; **Figure 3B**), indicating a 10.0% decline in relation to model F. Model D showed the second lowest performance, with a PM_10 of 29.898, corresponding to a 7.2% reduction compared to model F. More details of the prediction performance for this data set are given in Table A2, Appendix A.

### Wheat_1 dataset

**Table 5** shows that, across all models, the G×E component explained the largest share of the variance, followed by the genotype component. Among the G×E subcomponents (VGE, VGEC, and VGEP), models C, D, E, and F captured much greater proportions of the total variability compared to model B, which included only the conventional G×E interaction term. Specifically, the increases relative to model B were approximately 83% for model C, 220% for model D, 379% for model E, and 467% for model F.

**Table 5.**
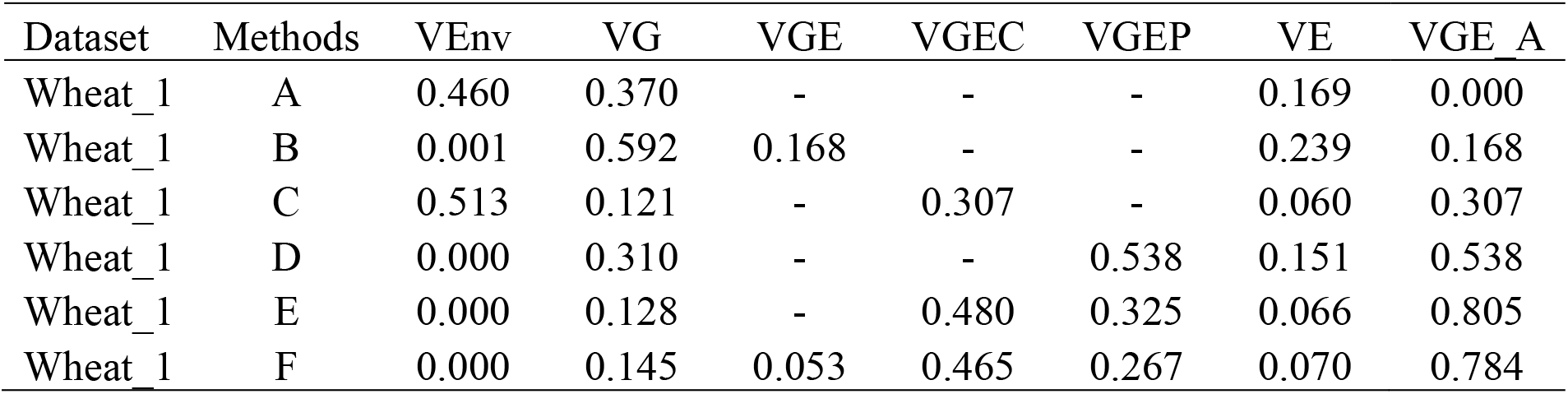
Proportion of total variances, in data set Wheat_1, explained for each variance component in each of the evaluated models. VGE_A = VGE + VGEC + VGEP, which denotes the total proportion of variance explained by the G×E interaction. 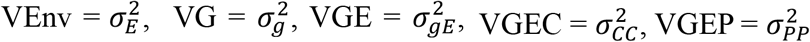, and VE = *σ*^2^.

| Dataset | Methods | VEnv | VG | VGE | VGEC | VGEP | VE | VGE A |
| --- | --- | --- | --- | --- | --- | --- | --- | --- |
| Wheat_1 | A | 0.460 | 0.370 | - | - | - | 0.169 | 0.000 |
| Wheat_1 | B | 0.001 | 0.592 | 0.168 | - | - | 0.239 | 0.168 |
| Wheat_1 | C | 0.513 | 0.121 | - | 0.307 | - | 0.060 | 0.307 |
| Wheat_1 | D | 0.000 | 0.310 | - | - | 0.538 | 0.151 | 0.538 |
| Wheat_1 | E | 0.000 | 0.128 | - | 0.480 | 0.325 | 0.066 | 0.805 |
| Wheat_1 | F | 0.000 | 0.145 | 0.053 | 0.465 | 0.267 | 0.070 | 0.784 |

**Table 6.** Proportion of total variance, across traits and data sets, explained for each variance component in each of the evaluated models. VGE_A = VGE + VGEC + VGEP, which denotes the total proportion of variance explained by the G×E interaction. 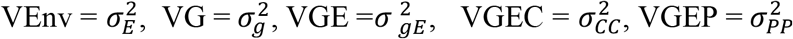, and VE = *σ*^2^. Time reports the execution time of each model across traits and data sets.

| Dataset | Methods | VEnv | VG | VGE | VGEC | VGEP | Ve | VGE A | Time |
| --- | --- | --- | --- | --- | --- | --- | --- | --- | --- |
| Across_data | A | 0.504 | 0.285 | - | - | - | 0.211 | 0.000 | 96.520 |
| Across_data | B | 0.402 | 0.318 | 0.109 | - | - | 0.171 | 0.109 | 331.640 |
| Across_data | C | 0.486 | 0.244 | - | 0.113 | - | 0.157 | 0.113 | 277.942 |
| Across_data | D | 0.432 | 0.255 | - | - | 0.154 | 0.160 | 0.154 | 268.559 |
| Across_data | E | 0.433 | 0.238 | - | 0.099 | 0.106 | 0.124 | 0.204 | 414.080 |
| Across_data | F | 0.401 | 0.257 | 0.041 | 0.099 | 0.101 | 0.100 | 0.242 | 638.115 |

Regarding COR, model F demonstrated the highest predictive performance (COR = 0.530), followed closely by model E (COR = 0.528), which was 0.4% lower than model F (Figure 4A). In contrast, model A showed the weakest performance (COR = 0.447; **Figure 4A**), representing an 18.6% decline relative to model F. Model B ranked second lowest, with a COR of 0.457, reflecting a 16% reduction compared to model F.

**Figure 4.**
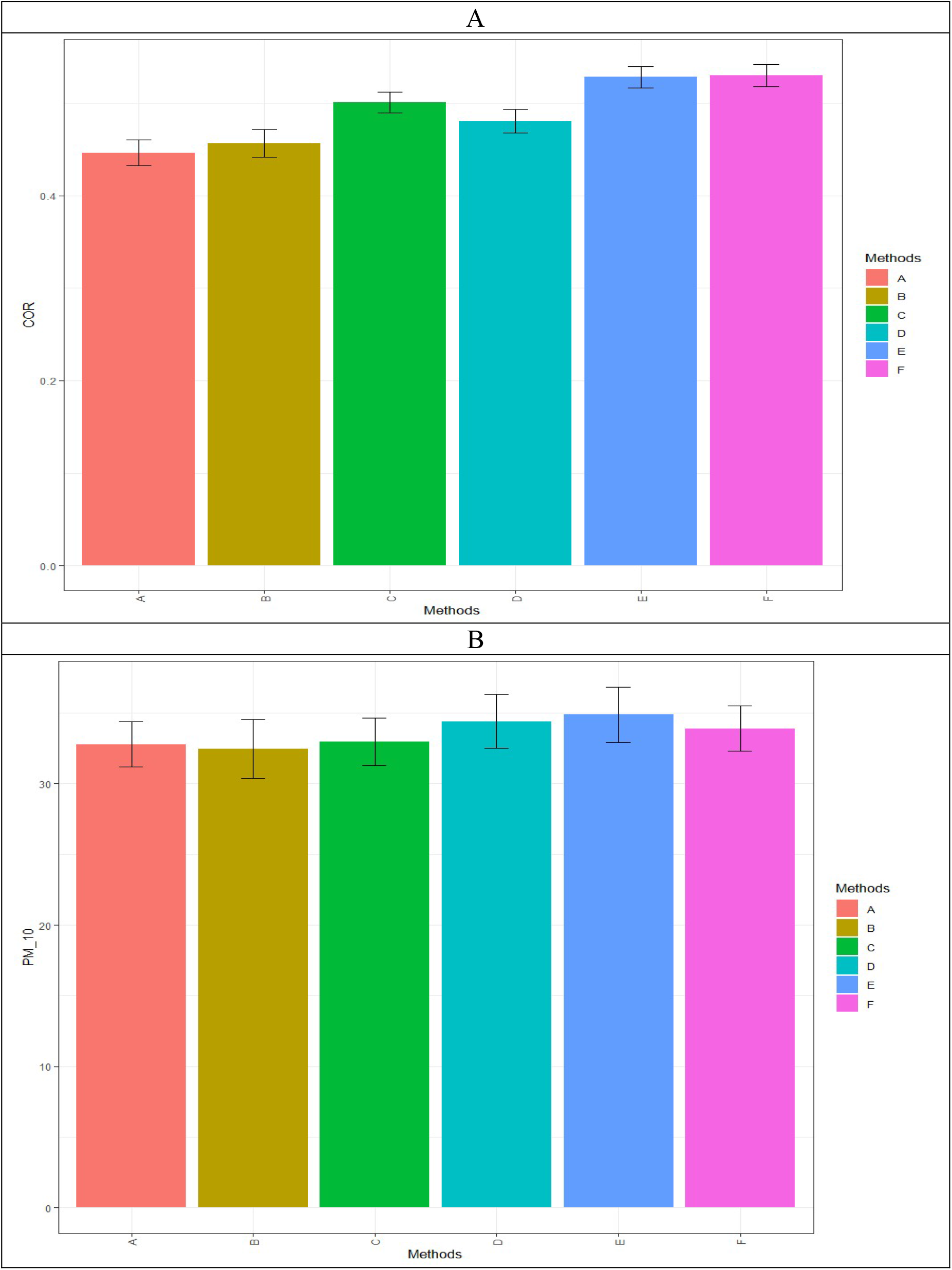
Prediction performance of the six evaluated methods on the Wheat_1 dataset, measured by average Pearson’s correlation (COR; A) and the proportion of test items whose predicted top 10% lines match the true top 10% (PM_10, B).

For PM_10, model E attained the best predictive outcome (PM_10 = 34.879), followed by model D (PM_10 = 34.411), which was 1.4% lower than model E (**Figure 4B**). Model B had the lowest result (PM_10 = 32.452; Figure 4B), a 7.5% decrease compared to model E, while model A exhibited the second lowest (PM_10 = 32.787); 6.4% lower than model E. More details of the prediction performance for this data set are given in Table A2, Appendix A.

### Across datasets

Across the 11datasets analyzed, **Table 6** shows that the environment component consistently explained the largest proportion of variance, followed by the genotype component. For the G×E subcomponents (VGE, VGEC, and VGEP), models C, D, E, and F accounted for much higher shares of the total variability compared with model B, which included only the conventional G×E interaction term. Specifically, the gains relative to model B were approximately 3.7% for model C, 41% for model D, 87% for model E, and 122% for model F. Notably, the increased variance explained in models C and D did not result in longer execution times; these models required about 16% (model C) and 19% (model D) less time than model B. In contrast, models E and F had longer execution times, exceeding that of model B by about 25% and 92%, respectively. The proportion of total variance in datasets EYT_2, EYT_3, Wheat_2, Wheat_3, Wheat_4, Wheat_5, and Wheat_6 is provided in **Table A1 of Appendix A**.

For COR, model F exhibited the strongest predictive capability (COR = 0.438), followed by model E (COR = 0.431), which was 1.6% lower than that of model F (**Figure 5A**). Model A had the lowest performance (COR = 0.355; Figure 5A), a 23.4% decrease relative to model F, while Model B was the second least effective (COR = 0.387; 13.2% lower than model F).

PM_10, model F achieved the highest predictive value (PM_10 = 30.117), followed by model E (PM_10 = 29.331), which was 2.7% lower than that of model F (**Figure 5B**). Model A had the lowest performance (PM_10 = 25.574; **Figure 5B**), a 17.8% decline compared to model F, while model D was the second lowest performance (PM_10 = 27.157; 10.94% lower than model E).

More details of the prediction performance across datasets are given in Table A2, Appendix A. While the prediction performance of the remaining datasets (EYT_2, EYT_3, Wheat_2, Wheat_3, Wheat_4, Wheat_5, and Wheat_6) is provided in **Table B1, Appendix B**.

**Figure 5.**
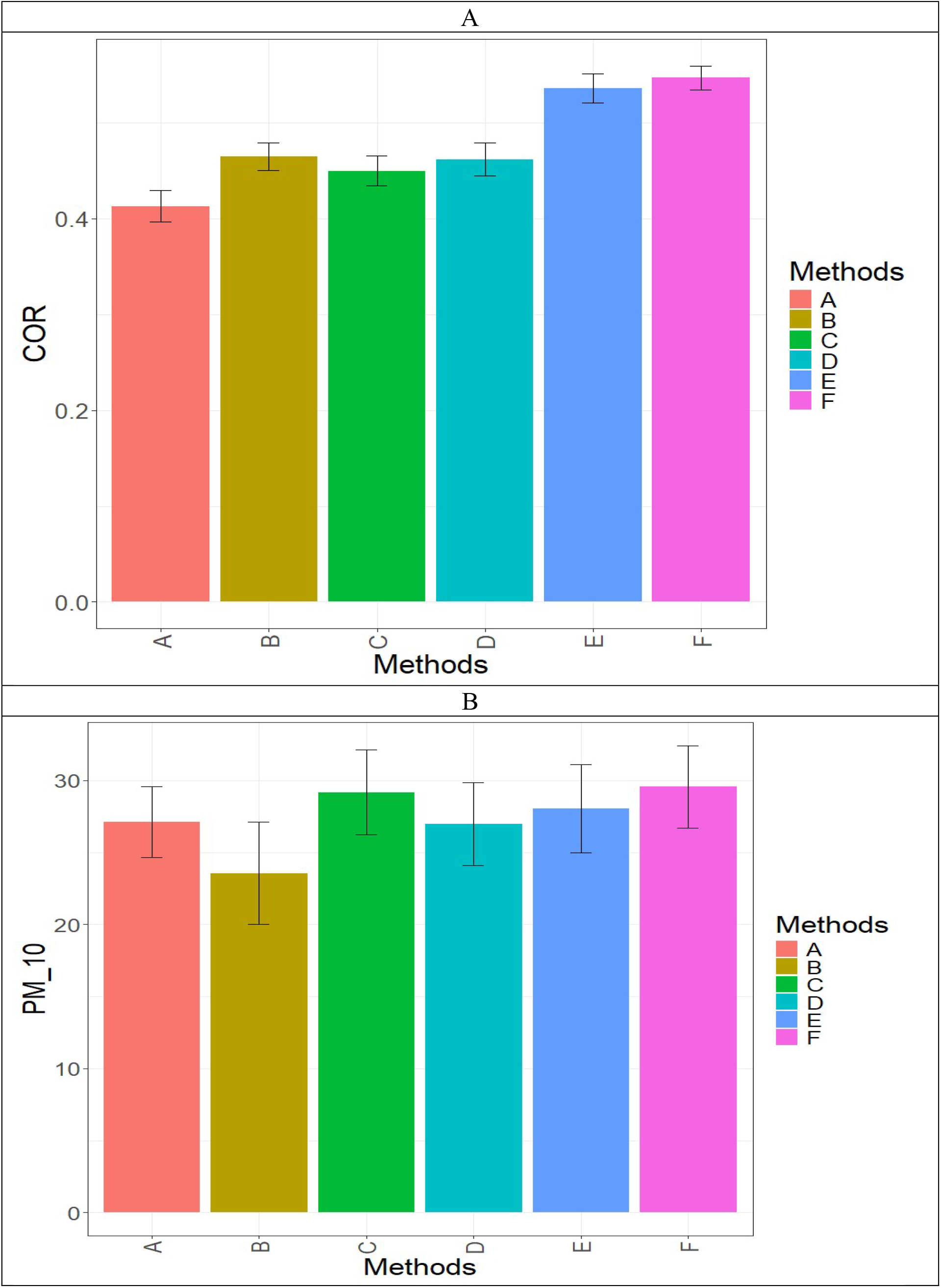
Prediction performance of the six evaluated methods across the dataset, measured by average Pearson’s correlation (COR; A) and the proportion of test items whose predicted top 10% lines match the true top 10% (PM_10, B).

## DISCUSSION

Modeling G×E interaction remains one of the most complex and critical challenges in the development of robust genomic prediction (GP) models. In plant breeding, G×E refers to the differential performance of genotypes across varying environmental conditions. Inadequate modeling of this interaction can lead to biased or inaccurate predictions, reducing the effectiveness of genomic selection in diverse agroecological systems (Crossa et al., 2017).

A central challenge in modeling G×E is capturing both environmental variability and the dynamic nature of genotype responses. Traditional GP models often treat G×E as random noise or as a simple additive component, which can oversimplify the interaction and fail to capture its structured variability, especially in multi-environment trials (Jarquín et al., 2014). To address this, models that incorporate covariance structures derived from genomic and environmental kernels have been proposed (Burgueño et al., 2012), with the Hadamard (elementwise) product being a widely used approach.

The Hadamard product captures G×E interaction through joint similarity between genotypes and environments, offering a biologically interpretable and statistically efficient framework that has been widely applied in genomic prediction. However, our results indicate that joint similarity alone does not fully capture G×E interaction, and that additional structured components contribute to the observed variability. The proposed framework complements, rather than replaces, the Hadamard-based formulation by capturing additional structure in G×E interaction, as demonstrated by the superior performance of the combined model.

To address these limitations, we proposed an alternative framework based on matrix multiplication between genomic and environmental covariance matrices, adapted from Cuevas et al. (2025). This approach yields two additional covariance structures derived from the upper and lower triangular components of the matrix product, which capture complementary aspects of G×E interaction. These components can be incorporated either as alternatives to, or in combination with, the conventional Hadamard-based interaction term, allowing for a more flexible representation of the interaction space.

Empirical validation using 11 independent datasets demonstrated the advantage of the proposed approach. When applied independently, the alternative framework (Model E) outperformed the conventional Hadamard-based model (Model B), with average improvements of 11.4% in Pearson’s correlation and 7.4% in PM_10_. More importantly, combining both approaches (Model F) led to further increases in prediction accuracy, with gains of approximately 13.2% in correlation and 10% in PM_10_ relative to the conventional model. These results indicate that the two frameworks capture complementary sources of variation and that their integration provides a more complete representation of G×E interaction.

The superior performance of Model F suggests that, in the datasets analyzed, G×E interaction may be multi-structured and not fully captured by a single covariance formulation. While the Hadamard product reflects joint similarity between genotypes and environments, the matrix-product-based approach extracts additional structured information embedded in the covariance relationships. Models C and D, which capture only partial aspects of this structure, showed inconsistent performance, further supporting the importance of integrating multiple components to adequately represent G×E.

The novelty of this work lies in adapting the framework of Cuevas et al. (2025), originally developed for integrating genomic and pedigree or multi-omics data, to the modeling of G×E interaction. This extension demonstrates that the structural advantages of matrix-product-based covariance constructions can be effectively leveraged to improve prediction accuracy in multi-environment trials. The consistent improvements observed across datasets highlight the robustness and general applicability of the proposed method.

Accurately modeling G×E interactions is essential for improving the predictive performance of genomic selection, as environmental factors play a major role in determining phenotypic expression and genotype adaptability. This challenge becomes even more critical under climate change, where increasing environmental variability and uncertainty reduce the effectiveness of traditional models (Costa-Neto et al., 2021; Elias et al., 2016). In these contexts, more flexible modeling strategies are necessary to capture heterogeneous and complex interaction patterns.

Our proposed framework contributes to addressing these challenges by providing a more expressive and flexible representation of G×E interaction. By incorporating covariance structures derived from matrix multiplication, the method captures structured variability beyond joint similarity, resulting in improved predictive performance. Although this approach may require additional computational resources, this is unlikely to be a limiting factor for most modern breeding programs.

In parallel, recent advances in machine learning and deep learning offer alternative approaches for modeling G×E through nonlinear and high-dimensional representations. While promising, these methods often face challenges related to interpretability, computational cost, and data requirements (Montesinos-López et al., 2021). In contrast, the framework proposed here maintains interpretability while extending the flexibility of conventional mixed models.

In summary, G×E interaction is a fundamental and complex component of genomic prediction that requires flexible and structured modeling approaches. The results of this study suggest that, for the datasets analyzed, combining conventional and matrix-product-based covariance structures may provide a more complete and effective representation of G×E interaction. This perspective opens new opportunities for developing more flexible and biologically meaningful covariance models for multi-environment genomic prediction in plant breeding.

The results of this study suggest that, for the datasets analyzed, combining conventional and matrix-product-based covariance structures may provide a more complete and effective representation of G×E interaction.

Importantly, the proposed framework is not a hybrid kernel approach, but rather an extension of matrix-product-based covariance constructions specifically for modeling G×E interaction.

## CONCLUSION

In this study, we propose an alternative framework for modeling G×E interaction, aimed at improving the prediction accuracy of genomic prediction models. Unlike conventional approaches, where the variance-covariance matrix of the interaction term is constructed using the Hadamard product of the genotype and environment variance-covariance matrices, our method is based on a subproduct derived from the matrix multiplication of these two matrices. Based on analyses across 11 datasets, our approach significantly outperforms the traditional Hadamard-based method in terms of prediction accuracy. To strengthen the empirical support for our framework, we encourage other researchers to evaluate its performance using other datasets.

## DECLARATIONS

### Author contribution statement

All authors contributed to the conception of the study. OAML developed the R program for implementing the proposed method. Data analysis and writing were performed by OAML, AML, JCML, JC, CMHS, SD and RO. Writing was revised and edited by all authors. All authors discussed and interpreted the results, read, and approved the final manuscript.

## Acknowledgements

Not applicable.

## Data availability

The datasets used in this study are publicly available at: https://github.com/osval78/New_GE_Framework. This repository includes the phenotypic and genomic data for all analyses, as well as the code used to implement the proposed methodology.

## APPENDIX THEORY

### Positive Semidefinite and Construction of Covariance Structures in Models A–F

This appendix provides the mathematical justification for the covariance structures used in Models A–F. Since all random effects are assumed to follow multivariate normal distributions, their associated covariance matrices must be **symmetric and positive semidefinite (**PSD) or behave as such in practice.

#### A. Genomic relationship matrix G (Model A)

The genomic relationship matrix is defined as:

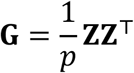

where **Z** is the standardized marker matrix. Because **G** is a Gram matrix, for any vector **x**:

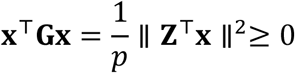

Thus, **G** is symmetric and positive semidefinite.

#### B. Environmental kernel K_*E*_

The environmental kernel is constructed as:

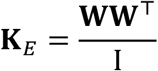

where **W** contains environmental covariates, or the design matrix of environments in the absence of environmental covariates. Thus, **K**_*E*_ is also a Gram matrix and is symmetric and PSD. It denotes the number of environments or the number of environmental covariates.

#### C. Genetic Kernel at the Observation Level: 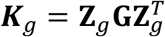

To show that 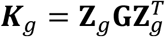 is PSD, we need to show that for any vector ***x, x***^*T*^ **K**_*g*_***x*** ≥ 0.

Let ***x*** be any real vector of appropriate dimension. Then:

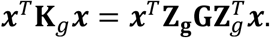

Define, 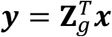, then the quadratic form becomes ***x***^*T*^**K**_*g*_ ***x*** = ***y***^*T*^**G*y***.

Since **G** is a genomic relationship matrix, and by construction it is PSD, therefore, for any vector ***y, y***^*T*^**G*y*** ≥ **0**. Thus

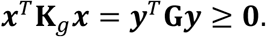

This holds for all vectors ***x***, so 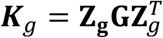 is PSD.

#### D. Hadamard Kernel K_*g*_*°*K_*E*_ (Model B)

The **Schur–Hadamard theorem** states:

If **A** and **B** are PSD matrices of the same dimension, then **A**°**B** is PSD.

Since both **K**_*g*_ and **K**_*E*_ are PSD, their Hadamard product **K**_*g*_*°***K**_*E*_ is also PSD. This ensures that the covariance matrix for the GxE interaction in Model B is valid.

#### E. Matrix-product-based kernels (Models C and D)

Let:

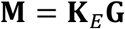

Although both **K**_*E*_and **G** are PSD, their matrix multiplication product **M** is **not necessarily symmetric nor** PSD. Define the upper-triangular operator as:

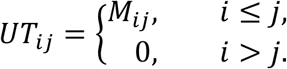

The diagonal operator is:

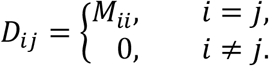

and the lower-triangular operator is:

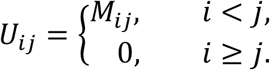

Thus, the full matrix, **M**, satisfies:

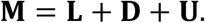

##### Model C (upper-triangular-based kernel)

The covariance matrix used in Model C is

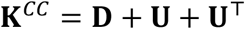

**D** is the diagonal part **U** is strictly upper triangular part. This matrix is symmetric by construction. Because **K**^*CC*^is obtained by symmetrizing a submatrix of **M**, and because symmetrization preserves PSD under the construction used by Cuevas et al. (2025), **K**^*CC*^ is PSD.

##### Model D (lower-triangular-based kernel)

Similarly, the covariance matrix in Model D is based on the **lower-triangular** component:

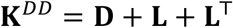

**L** is strictly lower triangular part.

###### Important remark

The matrices **K**^*CC*^and **K**^*DD*^are symmetric by construction. However, positive semidefiniteness is **not guaranteed analytically** for arbitrary matrix products.

Following the framework of Cuevas et al. (2025) and based on empirical evaluation in this study. Eigenvalue analysis confirmed that all constructed matrices had **non-negative spectra**. Furthermore, all matrices behaved as valid covariance structures across the 11 datasets analyzed.

#### F. Combined kernels (Models E and F)

##### Model E

Finally, the two components for the *G* × *E* for interaction are

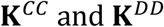

##### Model F

While for model F its three components for modeling the ***G*** × *E* for interaction are

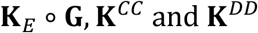

Since each component is a PSD matrix the resulting covariance matrices are valid for modeling.

#### G. Conceptual implication

The Hadamard kernel **K**_*E*_ ∘ **G** captures interaction through **joint similarity** between genotypes and environments.

In contrast, the matrix-product-based kernels **K**^*CC*^and **K**^*DD*^:

- extract **structured and directional information**
- represent **complementary aspects of G×E interaction**
- capture patterns not accessible through element-wise operations

This distinction provides a theoretical basis for the improved predictive performance observed in Models E and F.

##### Conclusion

All covariance matrices used in Models A–F are symmetric and behave as positive semidefinite in practice, ensuring that the corresponding multivariate normal models are well-defined. The proposed matrix-product-based constructions extend classical G×E modeling by introducing complementary covariance structures that enhance predictive performance.

## Appendix A

**Table A1.**
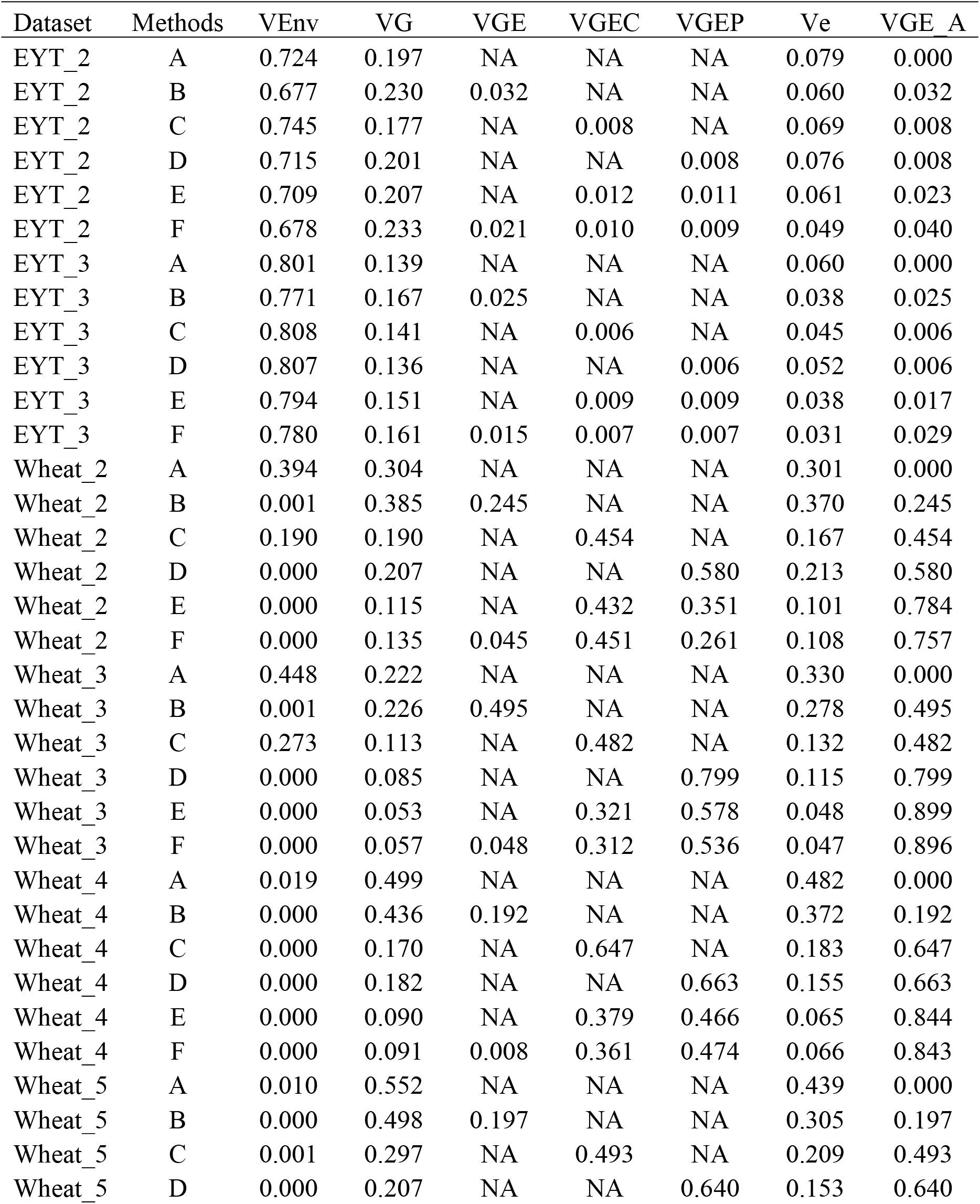
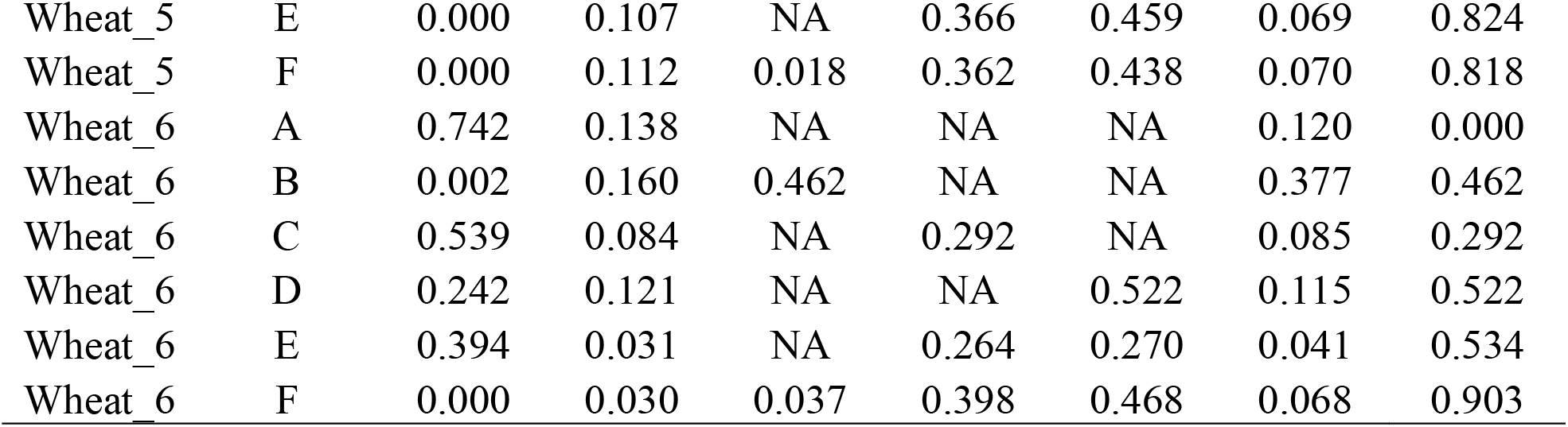
Proportion of total variance in data sets EYT_2, EYT_3, Wheat_2, Wheat_3, Wheat_4, Wheat_5, and Wheat_6, explained for each variance component in each of the evaluated models. VGE_A = VGE + VGEC + VGEP, which denotes the total proportion of variance explained by the G×E interaction. 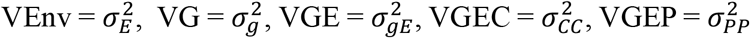, and VE = *σ*^2^.

| Dataset | Methods | VEnv | VG | VGE | VGEC | VGEP | Ve | VGE A |
| --- | --- | --- | --- | --- | --- | --- | --- | --- |
| EYT_2 | A | 0.724 | 0.197 | NA | NA | NA | 0.079 | 0.000 |
| EYT_2 | B | 0.677 | 0.230 | 0.032 | NA | NA | 0.060 | 0.032 |
| EYT_2 | C | 0.745 | 0.177 | NA | 0.008 | NA | 0.069 | 0.008 |
| EYT_2 | D | 0.715 | 0.201 | NA | NA | 0.008 | 0.076 | 0.008 |
| EYT_2 | E | 0.709 | 0.207 | NA | 0.012 | 0.011 | 0.061 | 0.023 |
| EYT_2 | F | 0.678 | 0.233 | 0.021 | 0.010 | 0.009 | 0.049 | 0.040 |
| EYT_3 | A | 0.801 | 0.139 | NA | NA | NA | 0.060 | 0.000 |
| EYT_3 | B | 0.771 | 0.167 | 0.025 | NA | NA | 0.038 | 0.025 |
| EYT_3 | C | 0.808 | 0.141 | NA | 0.006 | NA | 0.045 | 0.006 |
| EYT_3 | D | 0.807 | 0.136 | NA | NA | 0.006 | 0.052 | 0.006 |
| EYT_3 | E | 0.794 | 0.151 | NA | 0.009 | 0.009 | 0.038 | 0.017 |
| EYT_3 | F | 0.780 | 0.161 | 0.015 | 0.007 | 0.007 | 0.031 | 0.029 |
| Wheat_2 | A | 0.394 | 0.304 | NA | NA | NA | 0.301 | 0.000 |
| Wheat_2 | B | 0.001 | 0.385 | 0.245 | NA | NA | 0.370 | 0.245 |
| Wheat_2 | C | 0.190 | 0.190 | NA | 0.454 | NA | 0.167 | 0.454 |
| Wheat_2 | D | 0.000 | 0.207 | NA | NA | 0.580 | 0.213 | 0.580 |
| Wheat_2 | E | 0.000 | 0.115 | NA | 0.432 | 0.351 | 0.101 | 0.784 |
| Wheat_2 | F | 0.000 | 0.135 | 0.045 | 0.451 | 0.261 | 0.108 | 0.757 |
| Wheat_3 | A | 0.448 | 0.222 | NA | NA | NA | 0.330 | 0.000 |
| Wheat_3 | B | 0.001 | 0.226 | 0.495 | NA | NA | 0.278 | 0.495 |
| Wheat_3 | C | 0.273 | 0.113 | NA | 0.482 | NA | 0.132 | 0.482 |
| Wheat_3 | D | 0.000 | 0.085 | NA | NA | 0.799 | 0.115 | 0.799 |
| Wheat_3 | E | 0.000 | 0.053 | NA | 0.321 | 0.578 | 0.048 | 0.899 |
| Wheat_3 | F | 0.000 | 0.057 | 0.048 | 0.312 | 0.536 | 0.047 | 0.896 |
| Wheat_4 | A | 0.019 | 0.499 | NA | NA | NA | 0.482 | 0.000 |
| Wheat_4 | B | 0.000 | 0.436 | 0.192 | NA | NA | 0.372 | 0.192 |
| Wheat_4 | C | 0.000 | 0.170 | NA | 0.647 | NA | 0.183 | 0.647 |
| Wheat_4 | D | 0.000 | 0.182 | NA | NA | 0.663 | 0.155 | 0.663 |
| Wheat_4 | E | 0.000 | 0.090 | NA | 0.379 | 0.466 | 0.065 | 0.844 |
| Wheat_4 | F | 0.000 | 0.091 | 0.008 | 0.361 | 0.474 | 0.066 | 0.843 |
| Wheat_5 | A | 0.010 | 0.552 | NA | NA | NA | 0.439 | 0.000 |
| Wheat_5 | B | 0.000 | 0.498 | 0.197 | NA | NA | 0.305 | 0.197 |
| Wheat_5 | C | 0.001 | 0.297 | NA | 0.493 | NA | 0.209 | 0.493 |
| Wheat_5 | D | 0.000 | 0.207 | NA | NA | 0.640 | 0.153 | 0.640 |
| Wheat_5 | E | 0.000 | 0.107 | NA | 0.366 | 0.459 | 0.069 | 0.824 |
| Wheat_5 | F | 0.000 | 0.112 | 0.018 | 0.362 | 0.438 | 0.070 | 0.818 |
| Wheat_6 | A | 0.742 | 0.138 | NA | NA | NA | 0.120 | 0.000 |
| Wheat_6 | B | 0.002 | 0.160 | 0.462 | NA | NA | 0.377 | 0.462 |
| Wheat_6 | C | 0.539 | 0.084 | NA | 0.292 | NA | 0.085 | 0.292 |
| Wheat_6 | D | 0.242 | 0.121 | NA | NA | 0.522 | 0.115 | 0.522 |
| Wheat_6 | E | 0.394 | 0.031 | NA | 0.264 | 0.270 | 0.041 | 0.534 |
| Wheat_6 | F | 0.000 | 0.030 | 0.037 | 0.398 | 0.468 | 0.068 | 0.903 |

**Table A2.**
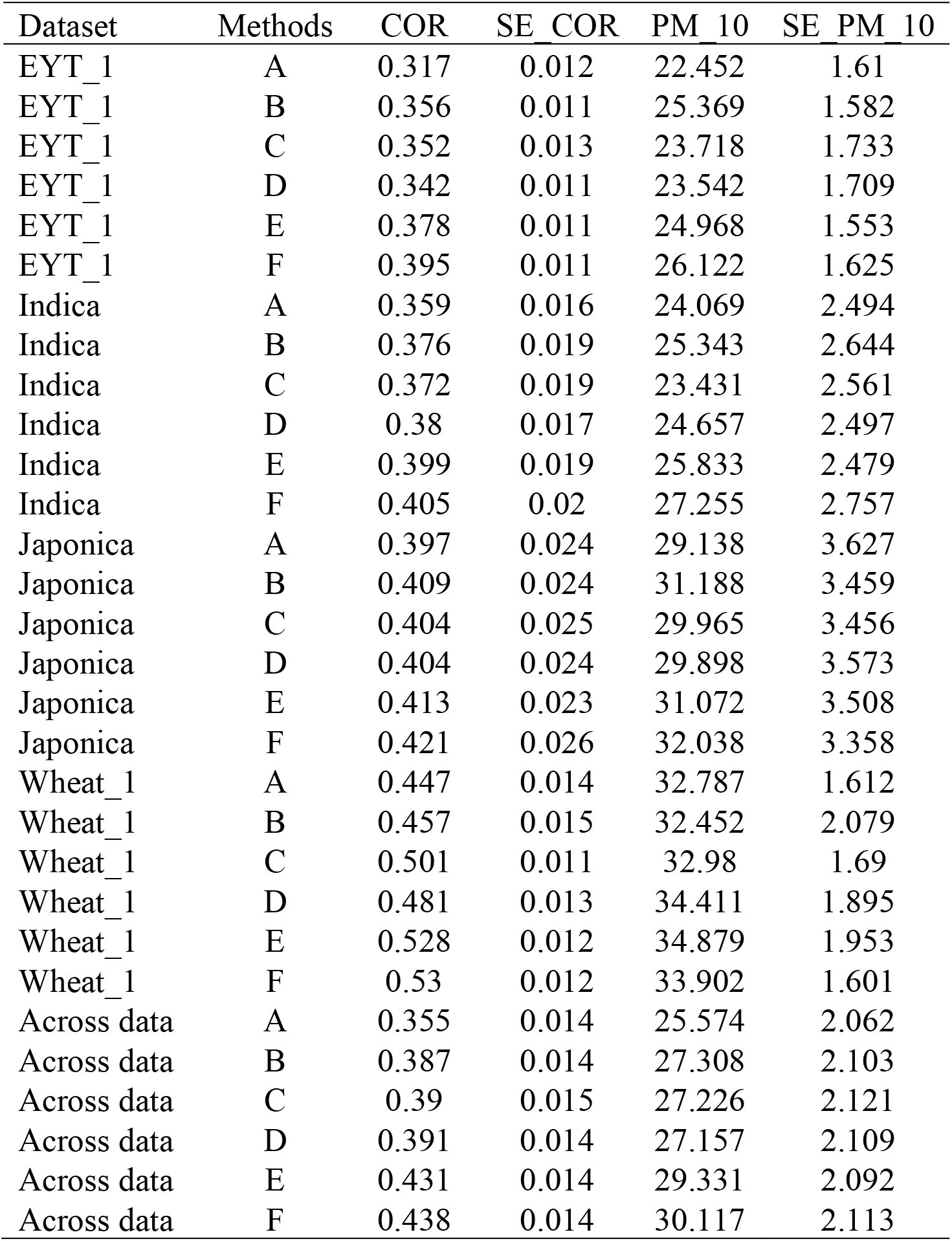
Prediction performance of the six evaluated methods on EYT_1 Indica, Japonica, Wheat_1 and across the dataset, measured by average Pearson’s correlation (COR) and the proportion of matching in the top 10% of lines (PM_10). SE_COR denotes the standard error for the COR metric, while SE_PM_10 is the standard error for the PM_10 metric.

## Appendix B

**Table B1.**
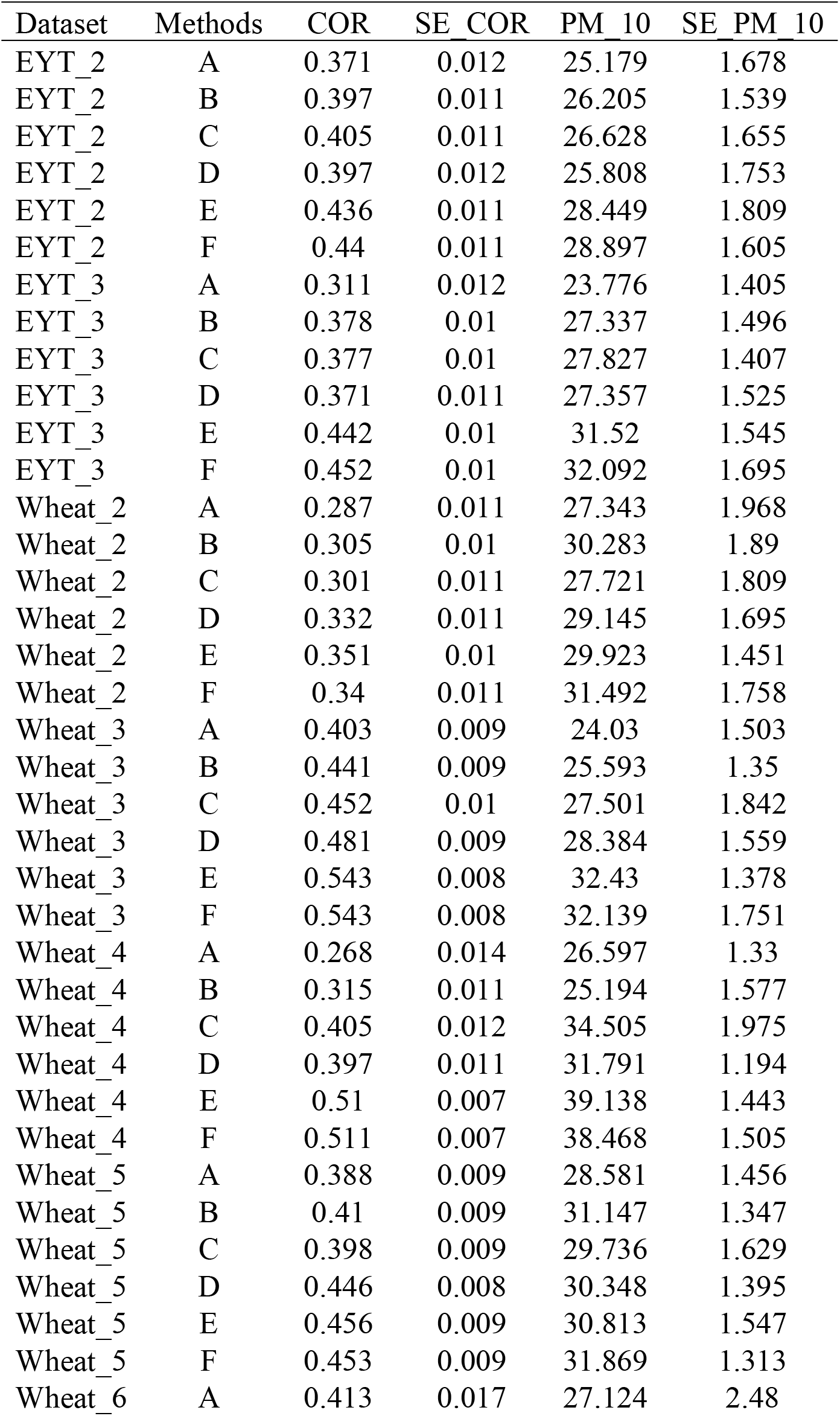
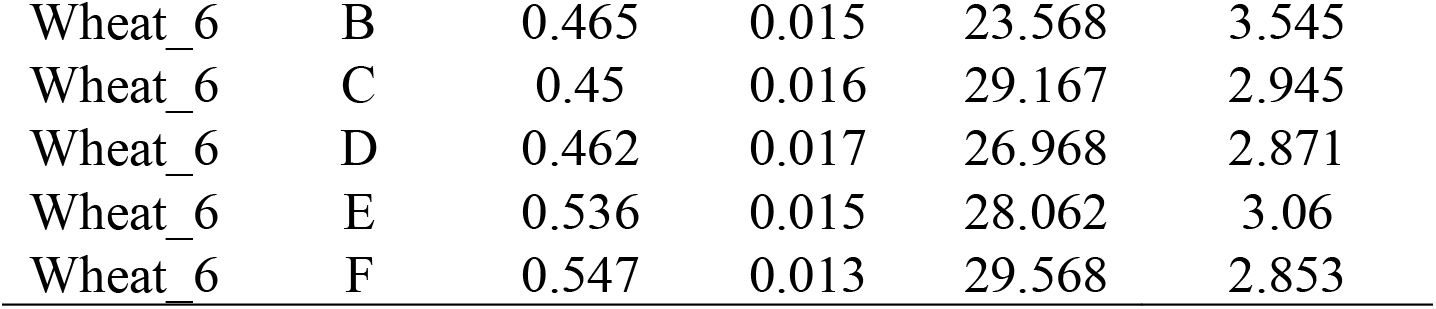
Prediction performance of the six evaluated methods on EYT_2, EYT_3, Wheat_2, Wheat_3, Wheat_4, Wheat_5, and Wheat_6 datasets, measured by average Pearson’s correlation (COR) and the proportion of matching in the top 10% of lines (PM_10). SE_COR denotes the standard error for the COR metric, while SE_PM_10 is the standard error for the PM_10 metric.

## REFERENCES

Alemu, A., Åstrand, J., Montesinos-López, O. A., Isidro Y Sánchez, J., Fernández-Gónzalez, J., Tadesse, W., Vetukuri, R. R., Carlsson, A. S., Ceplitis, A., Crossa, J., Ortiz, R., & Chawade, A. (2006). Genomic selection in plant breeding: Key factors shaping two decades of progress. Molecular Plant, 17(4), 552–578. 10.1016/j.molp.2024.03.007

Bradbury, P. J., Zhang, Z., Kroon, D. E., Casstevens, T. M., Ramdoss, Y., & Buckler, E. S. (2007). TASSEL: Software for association mapping of complex traits in diverse samples. Bioinformatics, 23(19), 2633–2635. 10.1093/bioinformatics/btm308

Burgueño, J., De Los Campos, G., Weigel, K., & Crossa, J. (2012). Genomic Prediction of Breeding Values when Modeling Genotype × Environment Interaction using Pedigree and Dense Molecular Markers. Crop Science, 52(2), 707–719. 10.2135/cropsci2011.06.0299

Cooper, M., Podlich, D., Jensen, N., Chapman, S., & Hammer, G. (1999). Modelling plant breeding programs. Trends in Agronomy, 2, 33–64.

Cooper, M., Technow, F., Messina, C., Gho, C., & Totir, L. R. (2016). Use of Crop Growth Models with Whole-Genome Prediction: Application to a Maize Multienvironment Trial. Crop Science, 56(5), 2141–2156. 10.2135/cropsci2015.08.0512

Costa-Neto, G., Fritsche-Neto, R., & Crossa, J. (2021). Nonlinear kernels, dominance, and envirotyping data increase the accuracy of genome-based prediction in multi-environment trials. Heredity, 126(1), 92–106. 10.1038/s41437-020-00353-1

Cornelius, P. L., Crossa, J., & Seyedsadr, M. S. (1992). Statistical tests and estimators of multiplicative models for genotype-by-environment interaction. Crop Science, 32, 193–198. 10.2135/cropsci1992.0011183X003200010043

Cornelius, P. L., & Crossa, J. (1999). Prediction assessment of shrinkage estimators of multiplicative models for multi-environment cultivar trials. Crop Science, 39, 998–1009.10.2135/cropsci1999.0011183X003900040012x

Crossa, J. (1990). Statistical analyses of multilocation trials. Advances in Agronomy, 44, 55–85.10.1016/S0065-2113(08)60818-4

Crossa, J., Campos, G. D. L., Pérez, P., Gianola, D., Burgueño, J., Araus, J. L., Makumbi, D., Singh, R. P., Dreisigacker, S., Yan, J., Arief, V., Banziger, M., & Braun, H.-J. (2010). Prediction of Genetic Values of Quantitative Traits in Plant Breeding Using Pedigree and Molecular Markers. Genetics, 186(2), 713–724. 10.1534/genetics.110.118521

Crossa, J., Fritsche-Neto, R., Montesinos-Lopez, O. A., Costa-Neto, G., Dreisigacker, S., Montesinos-Lopez, A., & Bentley, A. R. (2021). The Modern Plant Breeding Triangle: Optimizing the Use of Genomics, Phenomics, and Enviromics Data. Frontiers in Plant Science, 12, 651480. 10.3389/fpls.2021.651480

Crossa, J., Pérez-Rodríguez, P., Cuevas, J., Montesinos-López, O., Jarquín, D., De Los Campos, G., Burgueño, J., González-Camacho, J. M., Pérez-Elizalde, S., Beyene, Y., Dreisigacker, S., Singh, R., Zhang, X., Gowda, M., Roorkiwal, M., Rutkoski, J., & Varshney, R. K. (2017). Genomic Selection in Plant Breeding: Methods, Models, and Perspectives. Trends in Plant Science, 22(11), 961–975. 10.1016/j.tplants.2017.08.011

Cuevas, J., Crossa, J., Montesinos-López, A., Martini, J. W. R., Gerard, G. S., Ortegón, J., Dreisigacker, S., Govindan, V., Pérez-Rodríguez, P., Saint Pierre, C., Herrera, L. A. C., Montesinos-López, O. A., & Vitale, P. (2025). Enhancing wheat genomic prediction by a hybrid kernel approach. Frontiers in Plant Science, 16, 1605202. 10.3389/fpls.2025.1605202

Cuevas, J., Crossa, J., Montesinos-López, O. A., Burgueño, J., Pérez-Rodríguez, P., & De Los Campos, G. (2017). Bayesian Genomic Prediction with Genotype×Environment Interaction Kernel Models. G3 Genes|Genomes|Genetics, 7(1), 41–53. 10.1534/g3.116.035584

Eberhart, S. A., & Russell, W. A. (1966). Stability Parameters for Comparing Varieties^1^. Crop Science, 6(1), 36–40. 10.2135/cropsci1966.0011183X000600010011x

Elias, A. A., Robbins, K. R., Doerge, R. W., & Tuinstra, M. R. (2016). Half a Century of Studying Genotype × Environment Interactions in Plant Breeding Experiments. Crop Science, 56(5), 2090–2105. 10.2135/cropsci2015.01.0061

Elshire, R. J., Glaubitz, J. C., Sun, Q., Poland, J. A., Kawamoto, K., Buckler, E. S., & Mitchell, S. E. (2011). A Robust, Simple Genotyping-by-Sequencing (GBS) Approach for High Diversity Species. PLoS ONE, 6(5), e19379. 10.1371/journal.pone.0019379

Endelman, J. B. (2011). Ridge Regression and Other Kernels for Genomic Selection with R Package rrBLUP. The Plant Genome, 4(3), 250–255. 10.3835/plantgenome2011.08.0024

Finlay, K., & Wilkinson, G. (1963). The analysis of adaptation in a plant-breeding programme. Australian Journal of Agricultural Research, 14(6), 742–754. 10.1071/AR9630742

Fisher, Ronald Aylmer. (1925). Statistical methods for research workers.

Gauch, H. G. (1988). Model Selection and Validation for Yield Trials with Interaction. Biometrics, 44(3), 705. 10.2307/2531585

Henderson, Charles R and others. (1984). Applications of linear models in animal breeding (Vol. 462). University of Guelph Guelph.

Heslot, N., Akdemir, D., Sorrells, M. E., & Jannink, J.-L. (2014). Integrating environmental covariates and crop modeling into the genomic selection framework to predict genotype by environment interactions. Theoretical and Applied Genetics, 127(2), 463–480. 10.1007/s00122-013-2231-5

Ibba, M. I., Crossa, J., Montesinos-López, O. A., Montesinos-López, A., Juliana, P., Guzman, C., Delorean, E., Dreisigacker, S., & Poland, J. (2020). Genome-based prediction of multiple wheat quality traits in multiple years. The Plant Genome, 13(3), e20034. 10.1002/tpg2.20034

Jarquín, D., Crossa, J., Lacaze, X., Du Cheyron, P., Daucourt, J., Lorgeou, J., Piraux, F., Guerreiro, L., Pérez, P., Calus, M., Burgueño, J., & De Los Campos, G. (2014). A reaction norm model for genomic selection using high-dimensional genomic and environmental data. Theoretical and Applied Genetics, 127(3), 595–607. 10.1007/s00122-013-2243-1

Juliana, P., Singh, R. P., Poland, J., Mondal, S., Crossa, J., Montesinos-López, O. A., Dreisigacker, S., Pérez-Rodríguez, P., Huerta-Espino, J., Crespo-Herrera, L., & Govindan, V. (2018). Prospects and Challenges of Applied Genomic Selection—A New Paradigm in Breeding for Grain Yield in Bread Wheat. The Plant Genome, 11(3), 180017. 10.3835/plantgenome2018.03.0017

Messina, C. D., Technow, F., Tang, T., Totir, R., Gho, C., & Cooper, M. (2018). Leveraging biological insight and environmental variation to improve phenotypic prediction: Integrating crop growth models (CGM) with whole genome prediction (WGP). European Journal of Agronomy, 100, 151–162. 10.1016/j.eja.2018.01.007

Meuwissen, T. H. E., Hayes, B. J., & Goddard, M. E. (2001). Prediction of Total Genetic Value Using Genome-Wide Dense Marker Maps. Genetics, 157(4), 1819–1829. 10.1093/genetics/157.4.1819

Money, D., Gardner, K., Migicovsky, Z., Schwaninger, H., Zhong, G.-Y., & Myles, S. (2015). LinkImpute: Fast and Accurate Genotype Imputation for Nonmodel Organisms. G3 Genes|Genomes|Genetics, 5(11), 2383–2390. 10.1534/g3.115.021667

Montesinos-López, A., Montesinos-López, O. A., Crossa, J., Burgueño, J., Eskridge, K. M., Falconi-Castillo, E., He, X., Singh, P., & Cichy, K. (2016). Genomic Bayesian Prediction Model for Count Data with Genotype × Environment Interaction. G3 Genes|Genomes|Genetics, 6(5), 1165–1177. 10.1534/g3.116.028118

Montesinos-López, O. A., Montesinos-López, A., Crossa, J., Montesinos-López, J. C., Mota-Sanchez, D., Estrada-González, F., Gillberg, J., Singh, R., Mondal, S., & Juliana, P. (2018). Prediction of Multiple-Trait and Multiple-Environment Genomic Data Using Recommender Systems. G3 Genes|Genomes|Genetics, 8(1), 131–147. 10.1534/g3.117.300309

Montesinos-López, O. A., Montesinos-López, A., Mosqueda-González, B. A., Delgado-Enciso, I., Chavira-Flores, M., Crossa, J., Dreisigacker, S., Sun, J., & Ortiz, R. (2025a). Genomic prediction powered by multi-omics data. Frontiers in Genetics, 16, 1636438. 10.3389/fgene.2025.1636438

Montesinos-López, O. A., Martínez-Regalado, J. A., Murillo-Avalos, C. L., Herr, A. W., Montesinos-López, A., Delgado-Enciso, I., … & Carter, A. H. (2025b). Hybrid kernels integrating genomic and multispectral data improve wheat genomic prediction accuracy. The Plant Genome, 19(1), e70171.

Montesinos-López, O. A., Montesinos-López, A., Pérez-Rodríguez, P., Barrón-López, J. A., Martini, J. W. R., Fajardo-Flores, S. B., Gaytan-Lugo, L. S., Santana-Mancilla, P. C., & Crossa, J. (2021). A review of deep learning applications for genomic selection. BMC Genomics, 22(1), 19. 10.1186/s12864-020-07319-x

Monteverde, E., Gutierrez, L., Blanco, P., Pérez De Vida, F., Rosas, J. E., Bonnecarrère, V., Quero, G., & McCouch, S. (2019). Integrating Molecular Markers and Environmental Covariates To Interpret Genotype by Environment Interaction in Rice (Oryza sativa L.) Grown in Subtropical Areas. G3 Genes|Genomes|Genetics, 9(5), 1519–1531. 10.1534/g3.119.400064

Pérez, P., & De Los Campos, G. (2014). Genome-Wide Regression and Prediction with the BGLR Statistical Package. Genetics, 198(2), 483–495. 10.1534/genetics.114.164442

Poland, J. A., Brown, P. J., Sorrells, M. E., & Jannink, J.-L. (2012). Development of High-Density Genetic Maps for Barley and Wheat Using a Novel Two-Enzyme Genotyping-by-Sequencing Approach. PLoS ONE, 7(2), e32253. 10.1371/journal.pone.0032253

VanRaden, P. M. (2008). Efficient Methods to Compute Genomic Predictions. Journal of Dairy Science, 91(11), 4414–4423. 10.3168/jds.2007-0980

Washburn, J. D., Burch, M. B., & Franco, J. A. V. (2020). Predictive breeding for maize: Making use of molecular phenotypes, machine learning, and physiological crop models. Crop Science, 60(2), 622–638. 10.1002/csc2.20052

Yan, W., Hunt, L. A., Sheng, Q., & Szlavnics, Z. (2000). Cultivar Evaluation and Mega-Environment Investigation Based on the GGE Biplot. Crop Science, 40(3), 597–605. 10.2135/cropsci2000.403597x

